# Tuberculosis-associated IFN-I induces Siglec-1 on tunneling nanotubes and favors HIV-1 spread in macrophages

**DOI:** 10.1101/836155

**Authors:** Maeva Dupont, Shanti Souriant, Luciana Balboa, Thien-Phong Vu Manh, Karine Pingris, Stella Rousset, Céline Cougoule, Yoann Rombouts, Renaud Poincloux, Myriam Ben Neji, Carolina Allers, Deepak Kaushal, Marcelo J. Kuroda, Susana Benet, Javier Martinez-Picado, Nuria Izquierdo-Useros, Maria del Carmen Sasiain, Isabelle Maridonneau-Parini, Olivier Neyrolles, Christel Vérollet, Geanncarlo Lugo-Villarino

## Abstract

While tuberculosis (TB) is a risk factor in HIV-1-infected individuals, the mechanisms by which *Mycobacterium tuberculosis* worsens HIV-1 pathogenesis remain poorly understood. Recently, we showed that HIV-1 infection and spread are exacerbated in macrophages exposed to TB-associated microenvironments due to tunneling nanotube (TNT) formation. To identify molecular factors associated with TNT function, we performed a transcriptomic analysis in these macrophages, and revealed the up-regulation of the lectin receptor Siglec-1. We demonstrate Siglec-1 expression depends on TB-mediated production of type I interferon. In co-infected non-human primates, Siglec-1 is highly expressed by alveolar macrophages, whose abundance correlates with pathology and activation of the type I interferon/STAT1 pathway. Intriguingly, Siglec-1 expression localizes exclusively on microtubule-containing TNT that are long and carry HIV-1 cargo. Siglec-1 depletion in macrophages decreases TNT length, diminishes HIV-1 capture and cell-to-cell transfer, and abrogates TB-driven exacerbation of HIV-1 infection. Altogether, we uncover a deleterious role for Siglec-1 in TB-HIV-1 co-infection, and its localization on TNT opens new avenues to understand cell-to-cell viral spread.

## INTRODUCTION

Co-infection with *Mycobacterium tuberculosis* (Mtb) and the human immunodeficiency virus (HIV-1), the agents of tuberculosis (TB) and acquired immunodeficiency syndrome (AIDS), respectively, is a major health issue. Indeed, TB is the most common illness in HIV-1-infected individuals, about 55% of TB notified patients are also infected with HIV-1, and about a fifth of the TB death toll occurs in HIV-1 co-infected individuals (WHO health report 2017). Clinical studies evidence a synergy between these two pathogens, which is associated with a spectrum of aberration in immune function (Esmail et al., 2018). Yet, while progress has been made towards understanding how HIV-1 enhances Mtb growth and spread, the mechanisms by which Mtb exacerbates HIV-1 infection require further investigation (Bell and Noursadeghi, 2018; Deffur et al., 2013; Diedrich and Flynn, 2011).

Besides CD4^+^ T cells, macrophages are infected by HIV-1 in humans and by the simian immunodeficiency virus (SIV), the most closely related lentivirus to HIV, in non-human primates (NHP) (Cribbs et al., 2015; Rodrigues et al., 2017). Recently, using a humanized mouse model, macrophages were shown to sustain HIV-1 infection and replication, even in the absence of T cells (Honeycutt et al., 2017; Honeycutt et al., 2016). This is in line with several studies characterizing tissue macrophages, such as alveolar, microglia and gut macrophages, as reservoirs in HIV-1 patients undergoing antiretroviral therapy (Ganor et al., 2019; Jambo et al., 2014; Mathews et al., 2019; Sattentau and Stevenson, 2016).

Macrophages are key host cells for Mtb (O’Garra et al., 2013; VanderVen et al., 2016). We recently reported the importance of macrophages in HIV-1 exacerbation within the TB co-infection context (Souriant et al., 2019). Using relevant *in vitro* and *in vivo* models, we showed that TB-associated microenvironments activate macrophages towards an M(IL-10) profile, distinguished by a CD16^+^CD163^+^MerTK^+^ phenotype. Acquisition of this phenotype is dependent on the IL-10/STAT3 signaling pathway (Lastrucci et al., 2015). M(IL-10) macrophages are highly susceptible not only to Mtb infection (Lastrucci et al., 2015), but also to HIV-1 infection and spread (Souriant et al., 2019). At the functional level, we demonstrated that TB-associated microenvironments stimulate the formation of tunneling nanotubes (TNT), membranous channels connecting two or more cells over short to long distances above substrates. TNT are subdivided in two classes based on their thickness and cytoskeleton composition: “thin” TNT (<0.7 μm in diameter) containing F-actin, and “thick” TNT (>0.7 μm in diameter) are enriched in F-actin and microtubules (MTs) (Souriant et al., 2019). Thick TNT are functionally distinguished by the transfer of large organelles, such as lysosomes and mitochondria (Dupont et al., 2018; Eugenin et al., 2009; Hashimoto et al., 2016). While the contribution for each TNT class to HIV-1 pathogenesis has not been explored (Dupont et al., 2018; Eugenin et al., 2009; Hashimoto et al., 2016), we reported that total inhibition of TNT formation in M(IL-10) macrophages resulted in the abrogation of HIV-1 exacerbation induced by Mtb (Souriant et al., 2019). Factors influencing TNT function in M(IL-10) macrophages remain unknown at large.

In this study, global mapping of the M(IL-10) macrophage transcriptome revealed Siglec-1 (CD169, or sialoadhesin) as a potential factor responsible for HIV-1 dissemination in the co-infection context with TB. As a type I transmembrane lectin receptor, Siglec-1 possesses a large extracellular domain composed of 17 immunoglobulin-like domains, including the N-terminal V-set domain, which allows the *trans* recognition of terminal α2,3-linked sialic acid residues in *O*- and *N*-linked glycans and glycolipids, such as those surface-exposed in HIV-1 and SIV particles (Izquierdo-Useros et al., 2012a; Puryear et al., 2012). While Siglec-1 has yet to be implicated in the TB context, it is clearly involved in the pathogenesis of HIV-1, SIV and other retroviruses (Martinez-Picado et al., 2017). Siglec-1 is mainly expressed in myeloid cells (macrophages and dendritic cells) and participates in HIV-1 transfer from myeloid cells to T cells, as well as in the initiation of virus-containing compartment (VCC) formation in macrophages (Izquierdo-Useros et al., 2012a; Izquierdo-Useros et al., 2012b; Puryear et al., 2013; Puryear et al., 2012), and in the viral dissemination *in vivo* (Akiyama et al., 2017; Izquierdo-Useros et al., 2012a; Sewald et al., 2015). Indeed, HIV-1 and other retroviruses have evolved the capacity to hijack the immune surveillance and housekeeping immunoregulatory functions of Siglec-1 (Izquierdo-Useros et al., 2014; O’Neill et al., 2013). Here, we investigate how Siglec-1 expression is induced by TB, and the role it has in the capture and transfer of HIV-1 by TB-induced M(IL-10) macrophages, in particular in the context of TNT.

## RESULTS

### Tuberculosis-associated microenvironments induce Siglec-1 in macrophages

TB-induced M(IL-10) macrophages are highly susceptible to HIV-1 infection and spread (Souriant et al., 2019). To assess the global gene expression landscape in these cells, we performed a genome-wide transcriptome analysis (GEO submission GSE139511). To this end, we employed our published *in vitro* model (Lastrucci et al., 2015), which relies on the use of conditioned medium from either mock- (cmCTR) or Mtb-infected (cmMTB) human macrophages. As we described and observed before and herein, cmMTB-treated cells were positive for the M(IL-10) markers (CD16^+^CD163^+^MerTK^+^ and phosphorylated STAT3), and displayed a high rate of HIV-1 infection, as compared to those treated with cmCTR (Lastrucci et al., 2015). A distinct 60 gene-transcript signature was defined in cmMTB-treated cells, using a combination of the expression level, statistical filters and hierarchical clustering; 51 genes were up-regulated and 9 genes were down-regulated in cmMTB-compared to cmCTR-treated cells (Figure 1A). We compared expression data of cmMTB- and cmCTR-treated cells to public genesets available from MSigDB (Broad Institute) using the gene set enrichment analysis (GSEA) algorithm (Subramanian et al., 2005). As shown in Figure S1A, a significant fraction of genes that were up-regulated in response to interferon (IFN) type I (*e.g.* IFNα) and II (*i.e.* IFNγ), were also found, as a group, significantly up-regulated in cmMTB-treated cells in comparison to control samples (FDR q-value: < 10E-3). IFN-stimulated genes (ISG) usually exert antiviral activities (McNab et al., 2015; Schneider et al., 2014) and cannot be inferred as obvious candidates to facilitate HIV-1 infection. However, among this ISG signature, the up-regulation of Siglec-1 (7.4-fold, adjusted p value of 0.0162) in cmMTB-treated cells captured our attention due to its known role in HIV-1 pathogenesis (Izquierdo-Useros et al., 2014; O’Neill et al., 2013). We confirmed high Siglec-1 expression in cmMTB-treated macrophages at the mRNA (Figure 1B) and cell-surface protein (Figure 1C, Figure S1B) levels, which was superior to the level obtained in HIV-1-infected cells (Figure S1C). Particularly, cmMTB-treated macrophages displayed high density of Siglec-1 surface expression applying a quantitative FACS assay that determines the absolute number of Siglec-1 antibody binding sites per cell (Figure 1D).

**Figure 1.**
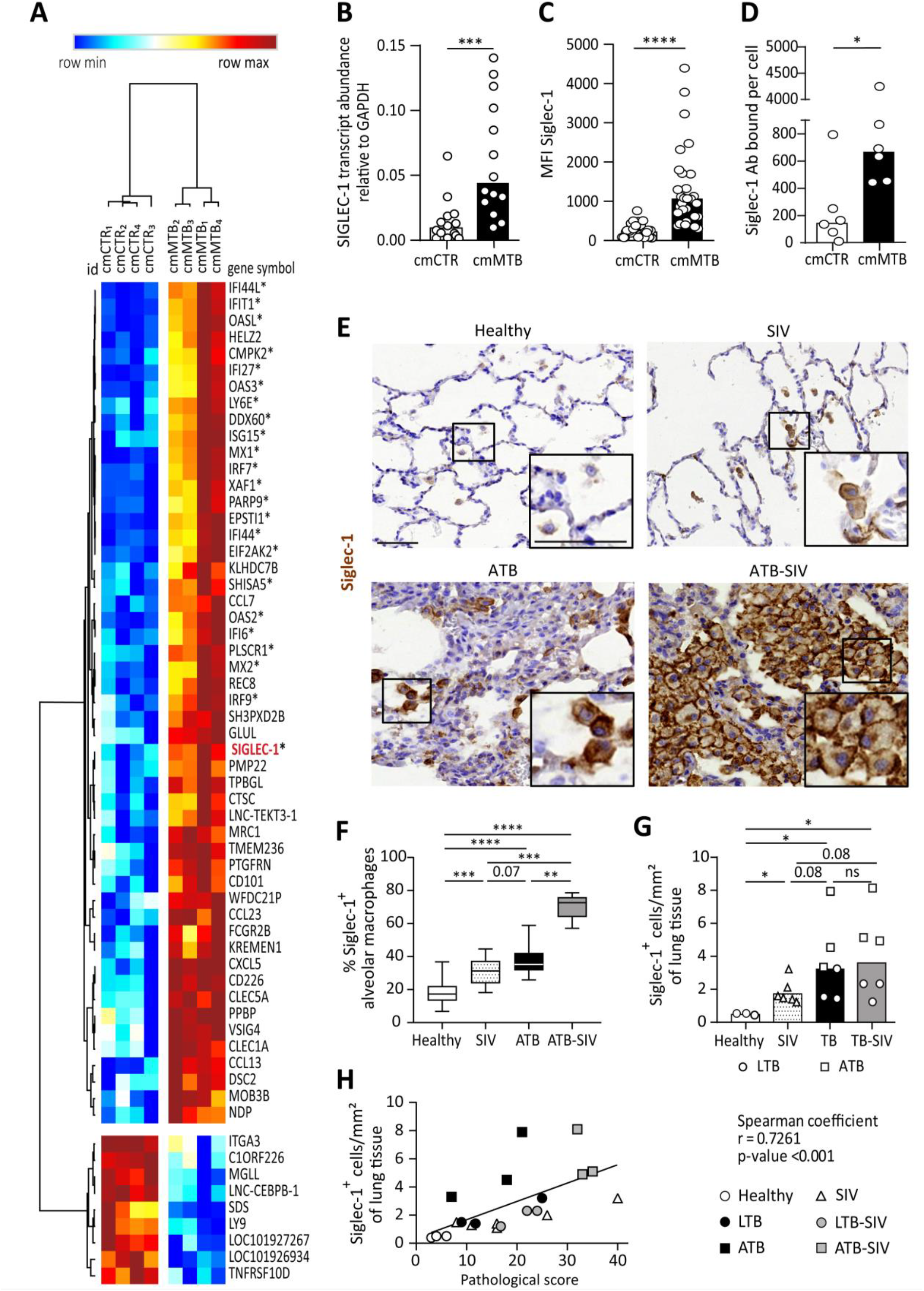
Tuberculosis-associated microenvironment induces Siglec-1 expression in macrophages (see also Figure S1 and S2). (A-D) For 3 days, human monocytes were differentiated into macrophages with cmCTR (white) or cmMTB (black) supernatants. (A) Heatmap from a transcriptomic analysis (GEO submission GSE139511) illustrating the top 60 differentially expressed genes (DEGs) between cmCTR- or cmMTB-cells. Selection of the top DEGs was performed using an adjusted p-value ≤ 0.05, a fold change of at least 2, and a minimal expression of 6 in a log_2_ scale. Hierarchical clustering was performed using the complete linkage method and the Pearson correlation metric with Morpheus (Broad Institute). Interferon-stimulated genes (ISG) are labelled with an asterisk and Siglec-1 is indicated in red. (B-D) Validation of Siglec-1 expression in cmMTB-treated macrophages. Vertical scatter plots showing the relative abundance to mRNA (B), median fluorescent intensity (MFI) (C), and mean number of Siglec-1 antibody binding sites per cell (D). Each circle represents a single donor and histograms median values. (E) Representative immunohistochemical images of Siglec-1 staining (brown) in lung biopsies of healthy, SIV infected (SIV), active TB (ATB), and co-infected (ATB-SIV) non-human primates (NHP). Scale bar, 100 μm. Insets are 2× zoom. (F) Vertical Box and Whisker plot indicating the distribution of the percentage of Siglec-1^+^ alveolar macrophages in lung biopsies from the indicated NHP groups. n=800 alveolar macrophages analyzed per condition from one representative animal per NHP group. (G) Vertical scatter plots displaying the number of cells that are positive for Siglec-1 per mm² of lung biopsies from the indicated NHP groups. Each symbol represents a single animal per NHP group. (H) Correlation between Siglec-1^+^ cells per mm² of lung tissue and the pathological score for healthy (white circle), SIV^+^ (white triangles), latent (black circle) or active (black square) TB, and SIV^+^ with latent (grey circle) or active (grey square) TB. Each symbol represents a single animal per NHP group. Mean value is represented as a black line. Statistical analyses: Two-tailed, Wilcoxon signed-rank test (B-D), Mann-Whitney unpaired test (F-G), Spearman correlation (H). *P < 0.05, **P < 0.01, ***P < 0.001, ****P < 0.0001. ns: not significant.

These data indicate that Siglec-1 is highly expressed in human macrophages exposed to TB-associated microenvironments and potentially in the context of TB-HIV co-infection.

### Siglec-1^+^ alveolar macrophage abundance correlates with pathology in co-infected primates

NHP has been an invaluable *in vivo* model to better understand the role of macrophages in SIV/HIV pathogenesis (Merino et al., 2017). Considering Siglec-1 binds sialylated lipids present in the envelop of HIV-1 and SIV (Izquierdo-Useros et al., 2012a; Puryear et al., 2012), we examined the presence of Siglec-1 positive alveolar macrophages in lung biopsies obtained from different NHP groups: (i) co-infected with Mtb (active or latent TB) and SIV, (ii) mono-infected with Mtb (active or latent TB), (iii) mono-infected with SIV, and (iv) healthy (Table S1, Figure S2A) (Cai et al., 2015; Kuroda et al., 2018; Souriant et al., 2019). Histological immuno-staining confirmed the presence of Siglec-1^+^ alveolar macrophages in the lungs of healthy NHP (Figure 1E-F and S2A), and revealed its significant increase in NHP mono-infected with either Mtb or SIV (Figure 1E-F and S2A). Strikingly, we noticed a massive abundance of these cells in co-infected NHP (Figure 1E-F and S2A). Concerning the overall abundance of Siglec-1^+^ leukocytes in lungs, we observed a significant increased in all infected NHP in comparison to healthy, with a higher tendency in active TB or co-infected NHP (Figure 1G and S2A). In fact, the number of Siglec-1^+^ leukocytes correlated positively with the severity of NHP pathology (Figure 1H, Table S2). Based on their cell morphology, localization in alveoli, and co-expression with the macrophage marker CD163 (Figure S2A-C), Siglec-1^+^ cells were identified as alveolar macrophages.

Collectively, these data show that Siglec-1 is up-regulated in alveolar macrophages in the context of a retroviral co-infection with active TB.

### Siglec-1 expression is dependent on TB-mediated type I IFN signaling

Siglec-1 is an ISG whose expression is induced by IFN-I in myeloid cells (Hartnell et al., 2001). In addition to viral infection, IFN-I is also induced in TB and known to mainly play a detrimental role (McNab et al., 2015; Moreira-Teixeira et al., 2018). Siglec-1 expression has not been described in the TB context or in co-infection with retroviruses such as SIV or HIV-1, therefore we assessed whether IFN-I stimulates Siglec-1 expression in TB-associated microenvironments. First, we found that cmMTB contains high amounts of IFN-I compared to cmCTR (Figure 2A). Next, we showed that recombinant IFN-β significantly increased Siglec-1 cell-surface expression in macrophages, close to the level induced by cmMTB (Figure 2B). Interestingly, we observed a modest, albeit significant, induction of Siglec-1 expression in cells treated with interleukin 10 (IL-10), a cytokine we have previously showed to be abundant in cmMTB (Lastrucci et al., 2015) and that renders macrophages highly susceptible to HIV-1 infection (Souriant et al., 2019). However, IL-10 depletion had no effect on Siglec-1 expression by cmMTB-treated cells (Figure 2C). By contrast, blocking the IFN-I receptor (IFNAR2) during cmMTB treatment fully abolished expression of Siglec-1 (Figure 2D), indicating that IFN-I is the responsible factor for Siglec-1 up-regulation in cmMTB-treated cells.

**Figure 2.**
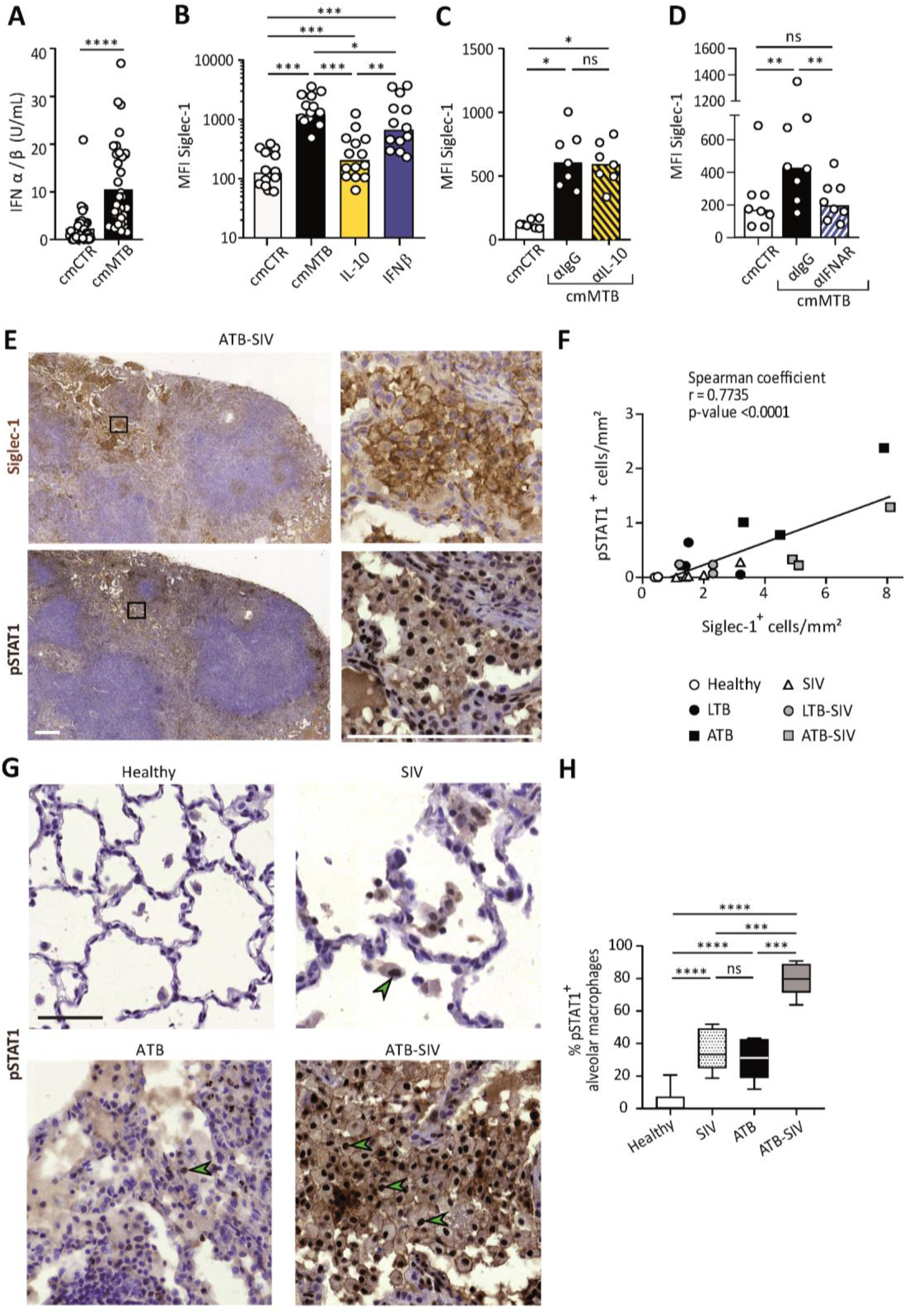
Siglec-1 expression is dependent on TB-mediated type I IFN signaling. (A) Vertical scatter plot showing the relative abundance of IFN-I in cmCTR (white) and cmMTB (black) media, as measured indirectly after 24h exposure to the HEK-Blue IFN-α/β reporter cell line yielding reporter activity in units (U) per mL. (B-D) For 3 days, monocytes were differentiated into macrophages either with cmMTB (black) or cmCTR (white), the indicated recombinant cytokines (B), the presence of an IL-10 depletion (α-IL-10) or a control (α-IgG) antibodies (C), or the presence of an IFNAR-2 blocking (α-IFNAR) or control (α-IgG) antibodies (D). (E) Representative serial immunohistochemical images of lung biopsies of a co-infected (ATB-SIV) NHP stained for Siglec-1 (brown, top) and pSTAT1 (brown, bottom). Scale bar, 250 μm. Insets are 10× zooms. (F) Correlation of the percentage of cells positive for Siglec-1 and pSTAT1, as measured per mm^2^ of lung tissue from the indicated NHP groups. Mean value is represented as a black line. (G) Representative immunohistochemical images of lung biopsies from the indicated NHP group stained for pSTAT1 (brown). Arrowheads show pSTAT1-positive nuclei. Scale bar, 500 μm. (H) Vertical Box and Whisker plot illustrating the percentage of pSTAT1^+^ alveolar macrophages in lung biopsies from the indicated NHP groups. N=450 alveolar macrophages analyzed per condition from one representative animal per NHP group. (A-D) Each circle within vertical scatter plots represents a single donor and histograms median value. (B-D) Vertical scatter plots displaying the median fluorescent intensity (MFI) of Siglec-1 cell-surface expression. Statistical analyses: Two-tailed, Wilcoxon signed-rank test (A-D), Spearman correlation (F), and Mann-Whitney unpaired test (H). *P < 0.05, **P < 0.01, ***P < 0.001, ****P < 0.0001. ns: not significant.

IFN-I binding to IFNAR leads to the phosphorylation and nuclear translocation of the transcription factor STAT1, whose role is essential for transcription of ISG (Ivashkiv and Donlin, 2014). We thus examined the status of STAT1 activation in co-infected NHP lung tissue. Histological staining of serial sections of co-infected lungs revealed that zones rich in Siglec-1^+^ leukocytes also exhibited positivity for nuclear phosphorylated STAT1 (pSTAT1) (Figure 2E), and the abundance of these two markers strongly correlated with the active status of TB in the different NHP groups (Figure 2F). Moreover, we found that the majority of Siglec-1^+^ alveolar macrophages were also positive for nuclear pSTAT1 in the infected NHP groups compared to healthy (Figure 2E and 2G). In fact, there was a higher number of pSTAT1^+^ alveolar macrophages in TB-SIV co-infected lungs when compared to those from mono-infected NHP (Figure 2G-H).

Altogether, these data demonstrate that Siglec-1 expression in human macrophages is controlled by IFN-I in a TB-associated microenvironment, and suggest the involvement of the IFN-I/STAT1/Siglec-1 axis in the pathogenesis of TB and co-infection with retroviruses.

### Siglec-1 localization on thick TNT is associated with their length and HIV-1 cargo

TNT formation is responsible for the increase in HIV-1 spread between human macrophages in TB-associated microenvironments (Souriant et al., 2019). To investigate whether Siglec-1 expression is involved in this process, we first examined its localization in the context of TNT formed by cmMTB-treated cells infected by HIV-1. We observed that Siglec-1 is localized exclusively on microtubule (MT)-positive thick TNT, and not on thin TNT (Figure 3A and Movie 1). Semi-automatic quantification of hundreds of TNT showed that about 50% of thick TNT were positive for Siglec-1 (Figure 3B and S3A). These TNT exhibited a greater length compared to those lacking Siglec-1 (Figure 3C). Importantly, unlike thin TNT, HIV-1 viral proteins are found mainly inside Siglec-1^+^ thick TNT (Figure 3D-E and Movie 2). In addition, these thick TNT also contain large organelles such as mitochondria (Figure 3F and S3B), another characteristic distinguishing thick from thin TNT (Dupont et al., 2018; Onfelt et al., 2006). In general, we also noticed that the incidence of Siglec-1^+^ thick TNT between HIV-1 infected macrophages persisted for more than one week upon HIV-1 infection (Figure S3C), suggesting a high degree of stability for these TNT.

**Figure 3.**
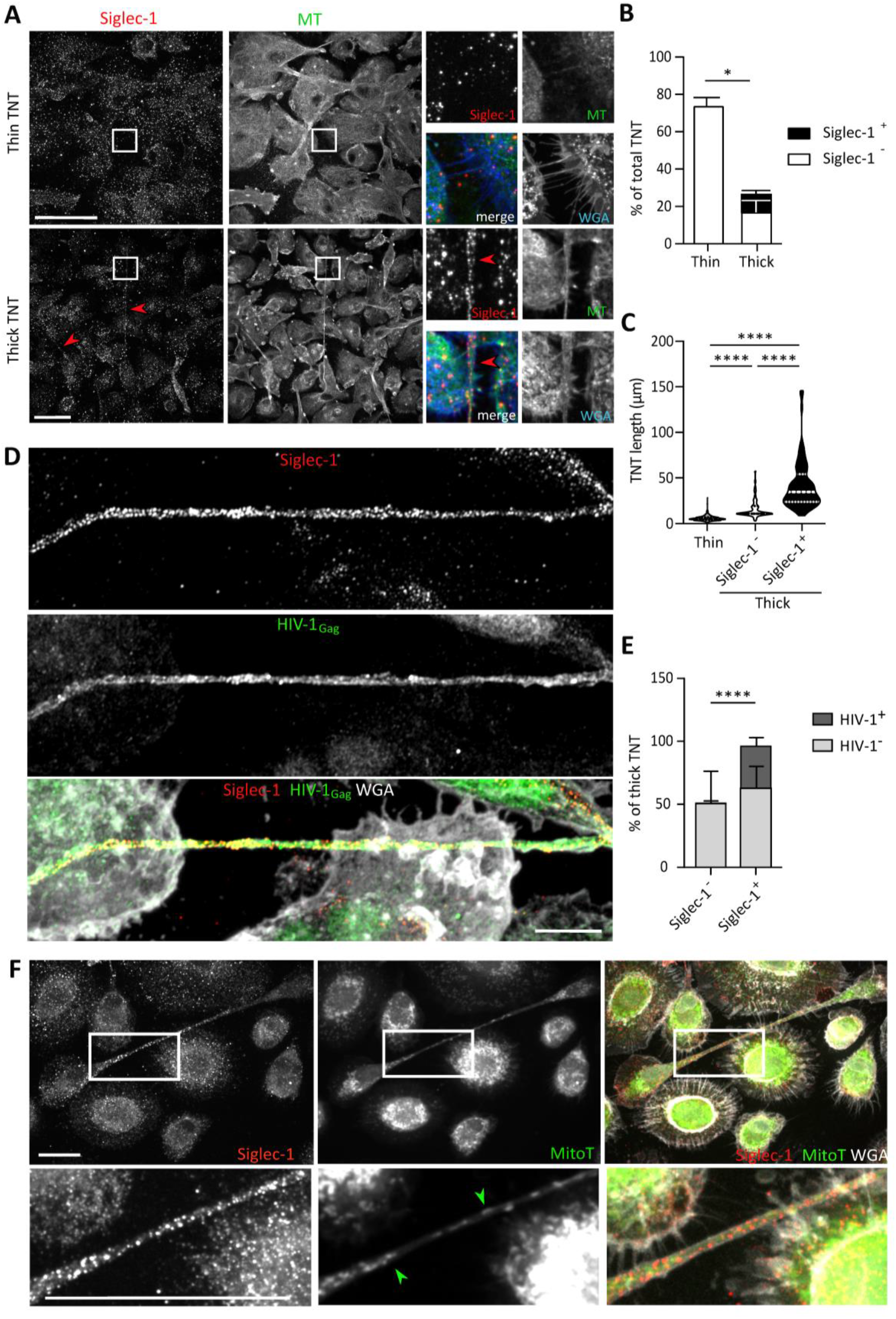
Siglec-1 localization on thick TNT is associated with their length and HIV/mitochondria cargo (see also Figure S3). (A-F) Human monocytes were differentiated into macrophages with cmMTB for 3 days, and then infected with HIV-1-ADA strain (unless indicated otherwise) and fixed 3 days post-infection. (A) Representative immunofluorescence images of cmMTB-treated macrophages infected with HIV-1-ADA, and stained for extracellular Siglec-1 (red), intracellular tubulin (MT, green) and Wheat Germ Agglutinin (WGA, blue). Inserts are 3× zooms. Red arrowheads show Siglec-1 localization on TNT. Scale bar, 20 μm. (B) Vertical bar plot showing the semi-automatic quantification of Siglec-1^+^ TNT (black) and Siglec-1^−^ TNT (white) in thick (WGA^+^, MT^+^) and thin (WGA^+^, MT^−^) TNT. 400 TNT were analyzed from 2 independent donors. (C) Siglec-1^+^ TNT exhibit a larger length index. Violin plots displaying the semi-automatic quantification of TNT length (in μm) for thin (WGA^+^, MT^−^), and thick TNT (WGA^+^, MT^+^) expressing Siglec-1 or not. 400 TNT were analyzed per condition from two independent donors. (D) Representative immunofluorescence images of cmMTB-treated macrophages 3-day post-infection with HIV-1-NLAD8-VSVG strain, and stained for extracellular Siglec-1 (red), intracellular HIV-1_Gag_ (green) and WGA (grey). Scale bar, 10 μm. (E) Vertical bar plots indicating the quantification of presence (dark grey) or absence (light grey) of HIV-1_Gag_ in thick TNT (WGA^+^, MT^+^) expressing Siglec-1 or not. 120 TNT in at least 1000 cells were analyzed from four independent donors. (F) Representative immunofluorescence images of cmMTB-treated macrophages infected with HIV-1-ADA loaded with MitoTracker (MitoT, green), and stained for extracellular Siglec-1 (red) and WGA (grey). Inserts are 3× zooms. Green arrowheads show mitochondria inside TNT. Scale bar, 10 μm. Statistical analyses: Two-way ANOVA comparing the presence of Siglec-1 in thin and thick TNT (B), and two-tailed Mann-Whitney unpaired test comparing TNT length (C) and the presence of HIV-1 in TNT (E). *P < 0.05, ****P < 0.0001.

These findings reveal an exclusive localization of Siglec-1 on MT-positive thick TNT that correlates positively with a greater length and high cargo of HIV-1 and mitochondria, arguing for a functional capacity of Siglec-1^+^ TNT to transfer material to recipient cells over long distances.

### TB-driven exacerbation of HIV-1 infection and spread in macrophages requires Siglec-1

To demonstrate a functional role for Siglec-1 in the susceptibility of macrophages to HIV-1 infection and spread induced by TB, Siglec-1 was depleted in cmMTB-treated cells by siRNA-mediated gene silencing (Figure 4A and S4A). While this depletion did not affect the total number of thick TNT (Figure 4B and S4B), we observed a 2-fold shortening of thick TNT in cells lacking Siglec-1 when compared to control cells (Figure 4C). Then, we performed a viral uptake assay in these cells using HIV-1 Gag-eGFP virus-like particles (GFP VLP) lacking the viral envelope glycoprotein but bearing sialylated lipids that interact with Siglec-1 on myeloid cells (Izquierdo-Useros et al., 2012b; Puryear et al., 2013). We consistently observed binding of VLP along Siglec-1^+^ thick TNT (Figure S4C). Yet, in the absence of Siglec-1, we noticed a significant reduction of VLP binding in comparison to control cells (Figure S4D). We confirmed this functional observation using a blocking monoclonal antibody against Siglec-1, showing that this receptor is involved in HIV-1 binding in cmMTB-treated cells (Figure S4E).

**Figure 4.**
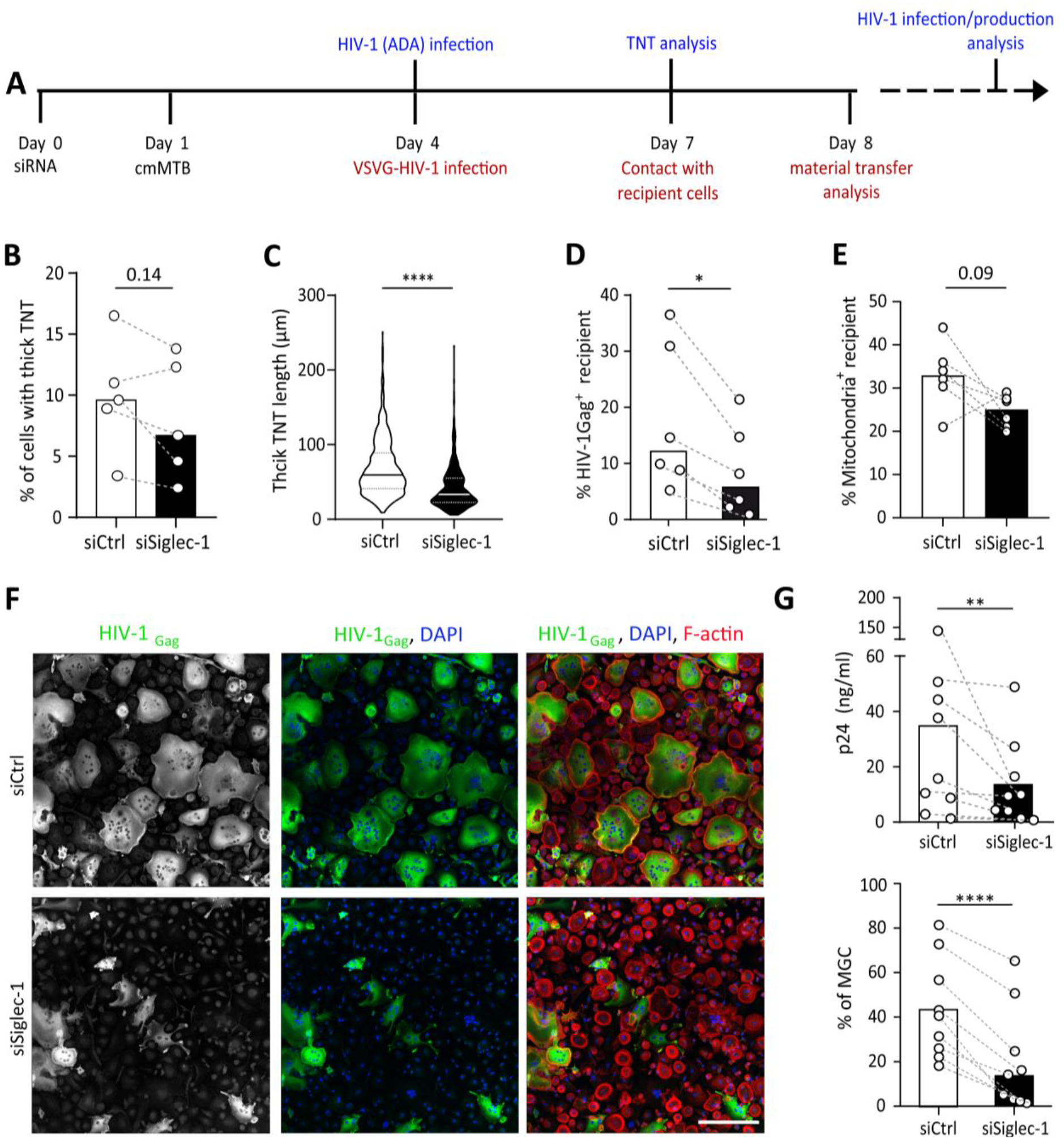
TB-driven exacerbation of HIV-1 infection and spread in macrophages requires Siglec-1 (see also Figure S4). (A) Experimental design. Monocytes from healthy subjects were transfected with siRNA targeting of Siglec-1 (siSiglec-1, black) or not (siCtrl, white). A day after, monocytes were differentiated into macrophages with cmMTB for 3 days. Cells were then infected with HIV-1-ADA (blue protocol) to measure the formation (B) and length (C) of TNT at day 7, or assess HIV-1 production and multinucleated giant cell (MGC) formation at day 14 (F-G). In parallel, cells were either infected with HIV-NLAD8-VSVG or labelled with mitoTracker to measure the transfer (red protocol) of HIV-1 (D) or mitochondria (E), respectively. (B) Before-and-after plots showing the percentage of cells forming thick TNT (F-actin^+^, WGA^+^, MT^+^). (C) Violin plots displaying the semi-automatic quantification of TNT length (in μm) for thick (WGA^+^, MT^+^) TNT; 300 TNT were analyzed per condition from two independent donors. (D-E) Before-and-after plots indicating the percentage of HIV-1_Gag_+ cells (D) or MitoTracker^+^ cells (E) among CellTracker^+^ cells after 24h co-culture. (F) Representative immunofluorescence images of siRNA transfected cells, treated with cmMTB 14 days post-HIV-1 infection. Cells were stained for intracellular HIV-1_Gag_ (green), Factin (red) and DAPI (blue). Scale bar, 500 μm. (G) Vertical scatter plots showing HIV-1-p24 concentration in cell supernatants (upper panel) and percentage of MGC (lower panel) at day 14 post-HIV-1 infection in cells represented in F (siSiglec-1, black; siCtrl, white). (B, D, E and G) Each circle represents a single donor and histograms median value. Statistical analyses: Paired t-test (B, G lower panel) or two-tailed, Wilcoxon signed-rank test (C-E, G upper panel). *P < 0.05, **P < 0.01, ****P < 0.0001.

We then assessed the role of Siglec-1 in HIV-1 transfer between macrophages, as this receptor is also important for the transfer of the virus to CD4^+^ T cells, (Akiyama et al., 2015; Izquierdo-Useros et al., 2012a; Puryear et al., 2013). We used an established co-culture system between cmMTB-treated macrophages that allows the transfer of the viral Gag protein from infected (donor, Gag^+^, red) to uninfected (recipient, CellTracker^+^, green) cells over 24 hours (Souriant et al., 2019) (Figure S4F). Of note, since Siglec-1 facilitates the infection of macrophages (Zou et al., 2011), we used VSV-G pseudotyped viruses to avoid any effect on HIV-1 primo-infection. Like this, we ensured the viral content was equal in cells at the time of the co-culture despite the loss of Siglec-1 (Figure S4G). The siRNA-mediated depletion of Siglec-1 significantly diminished the capacity of cmMTB-treated macrophages to transfer HIV-1 to recipient cells (Figure 4D), indicating that this receptor is involved in TB-driven macrophage-to-macrophage viral spread (Souriant et al., 2019). Intriguinly, there was a decreasing tendency for the capacity of Siglec-1-depleted cmMTB-treated macrophages to transfer mitochondria to recipient cells compared to controls (Figure 4E and S4F), alluding to a possible defect in mechanisms involved in intercellular material transfer including through thick TNT (Torralba et al., 2016). Remarkably, using replicative HIV-1 ADA strain (Figure 4A), we showed that silencing Siglec-1 expression in cmMTB-treated cells abolished the exacerbation of HIV-1 infection and production, as well as the enhanced formation of multinucleated giant cells (Figures 4F-G), which are pathological hallmarks of HIV-1 infection of macrophages (Verollet et al., 2015; Verollet et al., 2010).

These results determine that TB-induced Siglec-1 expression plays a key part in HIV-1 uptake and efficient cell-to-cell transfer, resulting in the exacerbation of HIV-1 infection and production in M(IL-10) macrophages.

## DISCUSSION

In this study, we investigated potential mechanisms by which Mtb exacerbates HIV-1 infection in macrophages, and uncovered a deleterious role for Siglec-1 in this process. These findings have different contributions to our understanding of this receptor in the synergy between Mtb and distinct retroviral infections, and also for TNT biology in host-pathogen interactions.

Our global transcriptomic approach revealed the up-regulation of Siglec-1, as part of an ISG-signature enhanced in macrophages exposed to a TB-associated microenvironment. Although pulmonary active TB has been characterized as an IFN-I-driven disease (Berry et al., 2010; McNab et al., 2015; Moreira-Teixeira et al., 2018), there are no reports in the literature about a role for Siglec-1 in TB or in Mtb co-infection with retroviruses. Expression of Siglec-1 is restricted to myeloid cells except circulating monocytes (Crocker et al., 2007), and is enhanced by IFN-I (Puryear et al., 2013; Rempel et al., 2008) and during HIV-1 infection (Pino et al., 2015). In addition, human alveolar macrophages are distinguished from lung interstitial macrophages by Siglec-1 expression (Yu et al., 2016). In this study, we determined that IFN-I present in TB-associated environments is responsible for Siglec-1 overexpression in human macrophages, which resembled that obtained in HIV-1-infected cells. While we saw a modest induction of Siglec-1 in macrophages upon IL-10 treatment, its depletion from the TB-associated microenvironment had no effect on Siglec-1 expression. This could be explained by the fact that IL-10 induces the autocrine production of IFN-I (Ziegler-Heitbrock et al., 2003) to indirectly modulate Siglec-1 expression in M(IL-10), which then contributes to the exacerbation of HIV-1 infection as we previously reported (Souriant et al., 2019). In the context of the most closely related lentivirus to HIV, namely SIV, we not only confirmed the presence of Siglec-1^+^ alveolar macrophages in SIV-infected NHP, but also reported the high abundance of these cells in active TB and in co-infected NHP groups, when compared to healthy ones. Importantly, we associated the high abundance of Siglec-1^+^ leukocytes with the increase NHP pathological scores, and it correlated positively to the detection of pSTAT1^+^ macrophage nuclei in histological staining of serial sections of lung biopsies from co-infected NHP. This is in line with a recent report on the presence of IFN-I, IFNAR and different ISGs in alveolar and lung interstitial tissue from NHP with active TB (Mattila, 2019), and with the fact that the *in vivo* expression of Siglec-1 is up-regulated early in myeloid cells after SIV infection and maintained thereafter in the pathogenic NHP model (Jaroenpool et al., 2007). In TB-SIV co-infection, we hypothesized that IFN-I is not exerting the expected antiviral effect, but instead is concomitant with chronic immune activation and attenuated by the high expression of Siglec-1 in myeloid cells, as recently proposed in the HIV-1 context (Akiyama et al., 2017). Altogether, these findings uncover the IFN-I/STAT1/Siglec-1 axis as a mechanism established by Mtb to exacerbate HIV-1 infection in myeloid cells, and call for the need to further investigate this signaling pathway in TB pathogenesis.

Another aspect worth highlighting is the impact that Siglec-1 expression has in the capture and transfer of HIV-1 by M(IL-10) macrophages, in particular in the context of TNT. First, we reported that Siglec-1 is located on MT-positive thick (and not on thin) TNT, correlating positively with increased length and HIV-1 cargo. To our knowledge, no receptor has been described so far to be present exclusively on thick TNT, making Siglec-1 an unprecedented potential marker for this subtype of TNT (Dupont et al., 2018). Second, viral uptake assays demonstrated the functional capacity of Siglec-1, including on thick TNT, to interact with viral-like particles bearing sialylated lipids; loss-of-function approaches showed Siglec-1 is important in the capture of these viral particles. Third, Siglec-1 depletion correlated with a decrease in thick TNT length, but had no effect in the total number of thick TNT. This suggests that, while the IFN-I/STAT1 axis is responsible for Siglec-1 expression in M(IL-10) macrophages, it does not contribute to TNT formation. This is line with our previous report where TNT formation induced by TB-associated microenvironments depends on the IL-10/STAT3 axis (Souriant et al., 2019). Concerning the shortening of thick TNT length, we infer that it may reflect a fragile state due to an altered cell membrane composition in the absence of Siglec-1; TNT are known for their fragility towards light exposure, shearing force and chemical fixation (Rustom et al., 2004). We hypothesize that the longer the TNT is, the more rigidity it requires to be stabilized. Cholesterol and lipids are known to increase membrane rigidity (Redondo-Morata et al., 2012) and are thought to be critical for TNT stability (Lokar et al., 2012; Thayanithy et al., 2014). Thus, the presence of Siglec-1 in thick TNT may affect the cholesterol and lipid composition *via* the recruitment of GM1/GM3 glycosphingolipid-enriched microvesicles (Halasz et al., 2018). In fact, TNT formation depends on GM1/GM3 ganglioside and cholesterol content (Kabaso et al., 2011; Lokar et al., 2012; Osteikoetxea-Molnar et al., 2016; Toth et al., 2017). Since GM1 and GM3 glycosphingolipids are *bona fide* ligands for Siglec-1 (Puryear et al., 2013), it is likely that Siglec-1^+^ thick TNT exhibit a higher lipid and cholesterol content, and hence an increase of membrane rigidity that favors the stability of longer TNT. Fourth, Siglec-1-depleted donor macrophages were less capable to transfer HIV-1, and to some extend mitochondria, to recipient cells. While infectious synapse and exososome release are mechanisms attributed to Siglec-1 that contribute to cell-to-cell transfer of HIV-1 (Bracq et al., 2018; Gummuluru et al., 2014; Izquierdo-Useros et al., 2014), they accomplish so extracellularly. Here, we speculate that Siglec-1 participates indirectly in the intracellular HIV-1 transfer *via* TNT as a tunnel over long distance, suggesting that factors affecting TNT rigidity favor distal viral dissemination while ensuring protection against immune detection. Finally, the depletion of Siglec-1 abrogated the exacerbation of HIV-1 infection and production induced by TB in M(IL-10) macrophages. This is likely to result from an accumulative effect of deficient capture and transfer of HIV-1 in the absence of Siglec-1. However, these results do not discern the specific contribution of Siglec-1 to the cell-to-cell transmission of HIV-1 *via* TNT from that obtained through other mechanisms (Bracq et al., 2018). Future studies will address whether the contribution of Siglec-1 to cell-to-cell transfer mechanisms has an impact in Mtb dissemination (Onfelt et al., 2006).

In conclusion, our study identifies Siglec-1 as a key TB-driven factor for the exacerbation of HIV-1 infection in macrophages, and as a new potential therapeutic target to limit viral dissemination in the co-infection context. It also sheds light on yet another housekeeping function for Siglec-1 with the potential to be hijacked by pathogens including HIV-1, such as intercellular communication facilitated by TNT. We argue that Siglec-1 localization on thick TNT has a physiological significance to macrophage biology in health and disease.

## MATERIAL AND METHODS

### Human Subjects

Monocytes from healthy subjects were provided by Etablissement Français du Sang (EFS), Toulouse, France, under contract 21/PLER/TOU/IPBS01/20130042. According to articles L12434 and R124361 of the French Public Health Code, the contract was approved by the French Ministry of Science and Technology (agreement number AC 2009921). Written informed consents were obtained from the donors before sample collection.

### Non-Human Primate (NHP) samples

All animal procedures were approved by the Institutional Animal Care and Use Committee of Tulane University, New Orleans, LA, and were performed in strict accordance with NIH guidelines. The twenty adult rhesus macaques used in this study (Table S1 and S2) were bred and housed at the Tulane National Primate Research Center (TNPRC). All macaques were infected as previously described (Foreman et al., 2016; Mehra et al., 2011; Souriant et al., 2019). Briefly, aerosol infection was performed on macaques using a low dose (25 CFU implanted) of Mtb CDC1551. Nine weeks later, a subgroup of the animals was additionally intravenously injected with 300 TCID50 of SIVmac239 in 1mL saline, while controls received an equal volume of saline solution. Euthanasia criteria were presentation of four or more of the following conditions: (i) body temperatures consistently greater than 2°F above pre-infection values for 3 or more weeks in a row; (ii) 15% or more loss in body weight; (iii) serum CRP values higher than 10 mg/mL for 3 or more consecutive weeks, CRP being a marker for systemic inflammation that exhibits a high degree of correlation with active TB in macaques (Kaushal et al., 2012; Mehra et al., 2011); (iv) CXR values higher than 2 on a scale of 0–4; (v) respiratory discomfort leading to vocalization; (vi) significant or complete loss of appetite; and (vii) detectable bacilli in BAL samples.

### Bacteria

Mtb H37Rv strain was grown in suspension in Middlebrook 7H9 medium (Difco) supplemented with 10% albumin-dextrose-catalase (ADC, Difco) and 0.05% Twen-80 (Sigma-Aldrich) (Lastrucci et al., 2015). For infection, growing Mtb was centrifuged (3000 RPM) at exponential phase stage and resuspended in PBS (MgCl_2_, CaCl_2_ free, Gibco). Twenty passages through a 26-G needle were done for dissociation of bacterial aggregates. Bacterial suspension concentration was then determined by measuring OD_600_, and then resuspended in RPMI-1640 containing 10% FBS for infection.

### Viruses

Virus stocks were generated by transient transfection of 293T cells with proviral plasmids coding for HIV-1 ADA and HIV-1 NLAD8-VSVG isolates, kindly provided by Serge Benichou (Institut Cochin, Paris, France), as previously described (Verollet et al., 2015). Supernatant were harvested 2 days post-transfection and HIV-1 p24 antigen concentration was assessed by a homemade enzyme-linked immunosorbent assay (ELISA). HIV-1 infectious units were quantified, as reported (Souriant et al., 2019) using TZM-bl cells (NIH AIDS Reagent Program, Division of AIDS, NIAID, NIH from Dr. John C. Kappes, Dr. Xiaoyun Wu and Tranzyme Inc).

HIV-VLP stock (GFP VLP) was generated by transfecting the molecular clone pGag-eGFP obtained from the NIH AIDS Research and Reference Reagent Program. HEK-293 T cells were transfected with calcium phosphate (CalPhos, Clontech) in T75 flasks using 30 μg of plasmid DNA. Supernatants containing VLPs were filtered (Millex HV, 0.45 μm; Millipore) and frozen at −80°C until use. The p24 Gag content of the VLPs was determined by an ELISA (Perkin-Elmer).

### Preparation of human monocytes and monocyte-derived macrophages

Human monocytes were isolated from healthy subject (HS) buffy coat (from EFS) and differentiated towards macrophages as described (Souriant et al., 2019). Briefly, peripheral blood mononuclear cells (PBMCs) were recovered by gradient centrifugation on Ficoll-Paque Plus (GE Healthcare). CD14^+^ monocytes were then isolated by positive selection magnetic sorting, using human CD14 Microbeads and LS columns (Miltenyi Biotec). Cells were then plated at 1.6 × 10^6^ cells per 6-well and allowed to differentiate for 5-7 days in RPMI-1640 medium (GIBCO), 10% Fetal Bovine Serum (FBS, Sigma-Aldrich) and human M-CSF (20 ng/mL) Peprotech) before infection with Mtb H37Rv for conditioned-media preparation. The cell medium was renewed every 3 or 4 days.

### Preparation of conditioned media

Conditioned-media from Mtb-infected macrophages (cmMTB) has been reported previously (Lastrucci et al., 2015; Souriant et al., 2019). Succinctly, hMDM were infected with Mtb H37Rv at a MOI of 3. After 18h of infection at 37°C, culture supernatants were collected, sterilized by double filtration (0.2μm pores) and aliquots were stored at −80°C. We then tested the capacity of individual cmMTB to differentiate freshly isolated CD14^+^ monocytes towards the M(IL-10) cell-surface marker phenotype, as assessed by FACS analyses. Those supernatants yielding a positive readout were then pooled together (5-10 donors) to minimize the inter-variability obtained between donors. Control media (cmCTR) was obtained from uninfected macrophages supernatant. When specified, IL-10 was eliminated from cmMTB by antibody depletion as described previously (Lastrucci et al., 2015; Souriant et al., 2019). The depletion was verified by ELISA (BD-Bioscience), according to manufacturer’s protocol.

### Conditioning of monocytes with the secretome of Mtb-infected macrophages or cytokines

Human CD14^+^ sorted monocytes from HS buffy coat were allowed to adhere in the absence of serum (0.4 × 10^6^ cells / 24-well in 500 μL) on glass coverslips, and then cultured for 3 days with 40% dilution (vol/vol) of cmCTR or cmMTB supplemented with 20% FBS and M-CSF (20 ng/mL, Peprotech). Blocking IFNAR receptor was performed by pre-incubation with mouse anti-IFNAR antibody (20 μg/mL, Thermo Fischer Scientific) in a 200 μL for 30 min prior to conditioning. After 3 days, cells were washed and collected for phenotyping.

When specified, monocytes were also conditioned in presence of 20 ng/mL M-CSF and 10 ng/mL recombinant human IL-10 (PeproTech) or 10 U/mL of IFNβ (Peprotech). Cell-surface expression of Siglec-1 was measured by flow cytometry using standard procedures detailed hereafter.

### RNA extraction and transcriptomic analysis

Cells conditioned with cmCTR and cmMTB supernatants (approximately 1.5 million cells) were treated with TRIzol Reagent (Invitrogen) and stored at −80°C. Total RNA was extracted from the TRIzol samples using the RNeasy mini kit (Qiagen). RNA amount and purity (absorbance at 260/280 nm) was measured with the Nanodrop ND-1000 apparatus (Thermo Scientific). According to the manufacturer’s protocol, complementary DNA was then reverse transcribed from 1 μg total RNA with Moloney murine leukemia virus reverse transcriptase (Invitrogen), using random hexamer oligonucleotides for priming. The microarray analysis was performed using the Agilent Human GE 4×44 v2 (single color), as previously described (Lugo-Villarino et al., 2018). Briefly, we performed hybridization with 2 μg Cy3-cDNA and the hybridization kit (Roche NimbleGen). The samples were then incubated for 5 min at 65°C, and 5 min at 42°C before loading for 17h at 42°C, according to manufacturer’s protocol. After washing, the microarrays were scanned with MS200 microarray scanner (Roche NimbleGen), and using Feature Extraction software, the Agilent raw files were extracted and then processed through Bioconductor (version 3.1) in the R statistical environment (version 3.6.0). Using the limma package, raw expression values were background corrected in a ‘normexp’ fashion and then quantile normalized and log_2_ transformed (Ritchie et al., 2015). Density plots, boxplots, principal component analyses, and hierarchical clustering assessed the quality of the hybridizations. Differentially expressed genes between macrophages exposed to cmCTR or cmMTB supernatants were extracted based on the p-value corrected using the Benjamini-Hochberg procedure. The log_2_ normalized expression values were used to perform Gene Set Enrichment Analyses (GSEA). The GSEA method allows to statistically test whether a set of genes of interest (referred to as a geneset) is distributed randomly or not in the list of genes that were pre-ranked according to their differential expression ratio between macrophages exposed to cmCTR or cmMTB supernatants. The output of GSEA is a GeneSet enrichment plot. The vertical black lines represent the projection onto the ranked GeneList of the individual genes of the GeneSet. The top curve in green corresponds to the calculation of the enrichment score (ES). The more the ES curve is shifted to the upper left of the graph, the more the GeneSet is enriched in the red cell population. Conversely, the more the ES curve is shifted to the lower right of the graph, the more the GeneSet is enriched in the blue cell population.

### siRNA silencing

Targets silencing in monocytes was performed using reverse transfection protocol as previously described (Troegeler et al., 2014). Shortly, human primary monocytes were transfected with 200 nM of ON-TARGETplus SMARTpool siRNA targeting Siglec-1 (Horizon Discovery) or non-targeting siRNA (control) using HiPerfect transfection system (Qiagen). Four hours post-transfection, transfected cells were incubated for 24h in RPMI-1640 medium, 10% FBS, 20 ng/mL of M-CSF, before addition of cmMTB media (40% vol/vol). After 3 additional days, cells were infected with HIV-1 ADA or HIV-1-NLAD8-VSV-G strain and kept in culture for 10 more days (48h, respectively). As soon as 3-day post-transfection, this protocol led to the efficient depletion of Siglec-1 between a range of 50-95%, as measured by flow cytometry.

### HIV-1 infection

For HIV-1 infection, at day 3 of differentiation, 0.4 × 10^6^ human monocytes-derived macrophages (hMDM) were infected with HIV-1 ADA strain (or specified) at MOI 0.1. HIV-1 infection and replication were assessed 10-day post-infection by measuring p24-positive cells by immunostaining and the level of p24 released in culture media by ELISA. For the infection and TNT quantification at day 6 post-infection, the same protocol was used. For HIV-1 transfer, higher MOI of HIV-1 VSVG pseudotyped NLAD8 virus was used, as described below (see section *HIV-1 and cell-to-cell transfer*) and in (Souriant et al., 2019).

### Uptake of Virus-Like Particles

Uptake experiment were performed as previously described (Izquierdo-Useros et al., 2012a; Izquierdo-Useros et al., 2014; Pino et al., 2015) using p24^Gag^ HIV-1_Gag−eGFP_ VLP (GFP VLP). Briefly, monocytes transfected or not with control siRNA or with siRNA directed against Siglec-1 and differentiated for 3 days in cmCTR or cmMTB were washed once with PBS prior to addition of 2 ng/mL of GFP VLP. Binding was performed during 3.5h at 37°C in a 5 % CO_2_ incubator. Cells were then detached with cell dissociation buffer (Gibco) and prepared for flow cytometry analysis on a BD LSRFortessa (TRI-Genotoul platerform). Same experiment was also performed blocking monocyte-derived macrophages at RT for 15 min with 10 μg/ml of mAb α-Siglec-1 7–239 (Abcam), IgG1 isotype control (BD Biosciences) or leaving cells untreated before VLP addition.

### Flow cytometry and Siglec-1 quantitation

Staining of conditioned macrophages was performed as previously described (Souriant et al., 2019). Adherent cells were harvested after 5 min incubation in trypsin 0.05% EDTA (Gibco) and washes with PBS (Gibco). After 10 min centrifugation at 320*g*, pellets were resuspended in cold staining buffer (PBS, 2mM EDTA, 0.5% FBS) with fluorophore-conjugated antibodies (See Table 1) and in parallel, with the corresponding isotype control antibody using a general concentration of 1 μg/mL. After staining, cells were washed with cold staining buffer, centrifuged for 2 min at 320*g* at 4°C, and analyzed by flow cytometry using BD LSRFortessa flow cytometer (BD Biosciences, TRI Genotoul plateform) and the associated BD FACSDiva software. Data were then analyzed using the FlowJo_V10 software (FlowJo, LLC). Gating on macrophage population was set according to its Forward Scatter (FSC) and Size Scatter (SSC) properties before doublet exclusion and analysis of the median fluorescence intensity (MFI) for each staining.

**Table 1.**

| REAGENT or RESOURCE | SOURCE | IDENTIFIER |
| --- | --- | --- |
| **Antibodies** |  |  |
| Mouse monoclonal anti-human Siglec-1 (clone 7-293) | Biolegend | Cat# 346008 |
| Mouse monoclonal anti-human CD16 (clone 3G8) | Biolegend | Cat# 302019 and 302018 |
| Mouse monoclonal anti-human CD163 (clone GHI/61) | Biolegend | Cat# 333608 |
| Mouse monoclonal anti-human MerTK (clone 590H11G1E3) | Biolegend | Cat# 367607 |
| Rabbit monoclonal anti-human STAT1 (clone 42H3) | Cell Signaling Technology | Cat# 9175 |
| Rabbit anti-human actin (a.a. 20-33) | Sigma-Aldrich | Cat# A5060 |
| Rabbit polyclonal anti-αtubulin | Abcam | Cat# ab18251 |
| Mouse monoclonal anti-Siglec-1 (clone hsn 7D2) | Novus Biologicals | Cat# NB 600-534 |
| Mouse monoclonal anti-Gag RD1 (clone KC57) | NIH AIDS Reagent program | Cat# 13449 |
| Monoclonal mouse anti-HIV p24 (clone Kal-1) | Dako Agilent technologies | Cat# M0857 |
| Mouse monoclonal anti-HIV-1 p24 (clone 183-H12-5C) | NIH AIDS Reagent Program | Cat# 3537 |
| Human polyclonal anti-HIV Immune Globulin (HIVIG) | NIH AIDS Reagent Program | Cat# 3957 |
| Polyclonal goat anti-human IgG | Sigma-Aldrich | Cat# A0170 |
| Mouse monoclonal anti-human CD163 (clone 10D6) | Leica/Novocastra | Cat# NCL-L-CD163 |
| Anti-pSTAT1 | Cell Signaling Technology | Cat# 9167 |
| Mouse monoclonal anti-IFNAR2 (clone MMHAR-2) | Thermo Fisher Scientific | Cat# 213851 |
| Mouse IgG2a isotype control | Thermo Fisher Scientific | Cat# 02-6200 |
| Polyclonal F(ab)2 goat anti-rabbit IgG, AlexaFluor 555 | Thermo Fisher Scientific | Cat# A-21430 |
| Polyclonal F(ab)2 goat anti-mouse IgG, AlexaFluor 488 | Thermo Fisher Scientific | Cat# A-10684 |
| Plyclonal F(ab)2 goat anti-mouse IgG, AlexaFluor 555 | Cell Signaling Technology | Cat# 4409 |
| Polyclonal goat anti-rabbit IgG, HRP | Thermo Fisher Scientific | Cat# 32460 |
| Polyclonal goat anti-mouse IgG, HRP | Thermo Fisher Scientific | Cat# 31430 |
| **Bacterial and Virus Strains** |  |  |
| *M. tuberculosis* H37Rv | N/A | N/A |
| HIV-1 ADA | Gift from Dr. S Benichou | Institut Cochin, Paris, France |
| HIV-1 ADA Gag-iGFP | Gift from Dr P. Benaroch and Dr. M. Schindler | Institu Pasteur, Paris, France<br>Institute of Virology, Munich, Germany |
| HIV-1 NLAD8-VSVG | Gift from Dr. S Benichou | Institut Cochin, Paris, France |
| HIV-1 ADA Gag-iGFP-VSVG | N/A | N/A |
| HIV-1 ADA-VSVG | Gift from Dr. S Benichou | Institut Cochin, Paris, France |
| HIV-1 Gag-eGFP | NIH research reagent program | Catalogue number 11468<br>Lot number: 2070514 |
| Biological Samples |  |  |
| Buffy Coat | Etablissement Français du Sang, Toulouse, France | N/A |
| Histological slides of lung biopsies from rhesus macaques | Tulane National Primate Research Center | N/A |
| Chemicals, Peptides, and Recombinant Proteins |  |  |
| Human recombinant M-CSF | Peprotech | Cat# 300-25 |
| Human recombinant IFNγ | Peprotech | Cat# 300-02BC |
| Human recombinant IL-10 | Peprotech | Cat# 200-10 |
| Critical Commercial Assays |  |  |
| Mouse anti-human CD14 microbeads | Miltenyi Biotec | Cat# 130-050-201 |
| LS magnetic columns | Miltenyi Biotec | Cat# 130-042-401 |
| Amersham ECL Prisme Western Blotting Detection Reagent | GE Healthcare | Cat# RPN2232 |
| SuperSignal WestPico Chemiluminescent Substrate | Thermo Scientific | Cat# 34080 |
| IL-10 ELISA set | BD Bioscience | Cat# 555157 |
| Trypsin EDTA 0.05% | Thermo Fisher Scientific | Cat# 25200072 |
| Accutase | Sigma-Aldrich | Cat# A-6964 |
| Phalloidin AlexaFluor 488 | Thermo Fisher Scientific | Cat# A12379 |
| Phalloidin Alexa Fluor 647 | Thermo Fisher Scientific | Cat# A22287 |
| DAPI | Sigma Aldrich | Cat# D9542 |
| CellTracker Red CMPTX Dye | Thermo Fisher Scientific | Cat# C34552 |
| CellTracker Green CMFDA Dye | Thermo Fisher Scientific | Cat# C7025 |
| MitoTracker Deep Red FM | Invitrogen | Cat# M22426 |
| Fluorescence Mounting Medium | Agilent Technologies | Cat# S302380-2 |
| Antibody diluent, Background reducing | DAKO, Agilent Technologies | Cat# S302283-2 |
| Experimental Models: Cell Lines |  |  |
| TZM-bl cell line | NIH AIDS Reagent Program | Cat# 8129 |
| HEK 293T cell line | NIH AIDS Reagent Program | Cat# 3318 |
| HKB-IFNAB | Invivogen | Cat# hb-detE |
| Software and Algorithms |  |  |
| ImageJ | ImageJ | http://www.imagej.nih.gov/ij |
| Prism (v8.0.0) | GraphPad | http://www.graphpad.com |
| Photoshop CS3 | Adobe | http://www.adobe.com |
| Adobe Illustrator CS5 | Adobe | https://www.adobe.com/fr/products/illustrator.html |
| Huygens Professional Version 16.10 | Scientific Volume Imaging | https://svi.nl/HuygensProfessional |
| FACS DIVA | BD Bioscience | http://www.bdbiosciences.com/ |
| FlowJo_v10 | FlowJo | https://www.flowjo.com/ |
| FCS Express V3 | DeNovo Software | http://www.denovosoftware.com |
| Image Lab | Bio-Rad Laboratories | http://www.bio-rad.com |
| Pannoramic Viewer | 3DHISTECH | https://www.3dhistech.com/pannoramic_viewer |

To determine Siglec-1 expression we applied a quantitative FACS assay. Briefly, cmCTR- and cmMTB-treated macrophages were detached using Accutase solution (Gibco) for 10 min at 37°C, washed, blocked with 1 mg/mL human IgG (Privigen, Behring CSL) and stained with mAb 7–239 α-Siglec-1-PE or matched isotype-PE control (Biolegend) at 4°C for 30 min. The mean number of Siglec-1 mAb binding sites per cell was obtained with a Quantibrite kit (Becton Dickinson) as previously described (Izquierdo-Useros et. al 2012b). Samples were analyzed with FACSCalibur using CellQuest software to evaluate collected data.

### Immunofluorescence microscopy

Cells were fixed with PFA 3.7%, Sucrose 30 mM in PBS. After washing with PBS, cells were saturated with blocking buffer (PBS-BSA 1%). Membrane proteins were then stained for 2h with primary antibodies: anti-Siglec-1 (10 μg/mL, Novus Biologicals). Cells were then incubated with appropriate secondary antibodies for 1h: Alexa Fluor 488 or 555 or 647 Goat anti-Mouse IgG (2 μg/mL, Cell Signaling Technology). Cells were then permeabilized as previously described (Souriant et al., 2019) with Triton X-100 0,3% for 10 minutes, washed in PBS before saturation with 0.6 μg/mL mouse IgG2 diluted in Dako Antibody Reducing Background buffer (Dako) for 30 min. Intracellular proteins were then stained with anti-Gag KC57 RD1 antibody (1/100, Beckman Coulter) and/or anti-α-tubulin (5 μg/mL, Abcam) for 2h. Cells were washed and finally incubated with Alexa Fluor 488, 555 or 647 Goat anti-Mouse or Goat anti-Rabbit IgG secondary antibodies (2 μg/mL, Cell Signaling Technology), Alexa Fluor 488 or 555 Phalloidin (33 mM, Thermo Fisher Scientific), Wheat Germ Agglutinin (CF^®^350 WGA, Thermofischer) and DAPI (500 ng/mL, Sigma Aldrich) in blocking buffer for 1h. Coverslips were mounted on a glass slide using Fluorescence Mounting Medium (Dako) and visualized with a spinning disk (Olympus) (Fig. S2C; 3A; 3F; S3A, S3B, S3C; movie 1), a Zeiss confocal LSM880 with Airyscan (Fig. 3D, Movie 2) and a FV1000 confocal microscope (Olympus) (Fig. 4F).

TNT were identified by WGA or phalloidin and tubulin staining, and counted on at least 1000 cells per condition and per donor.

As HIV-1 infection induces macrophages fusion into MGC (Verollet et al., 2010) the number of infected cells largely underestimates the rate of infection. Thus, to better reflect the rate of infection, we quantified the percentage of multinucleated cells. Using semi-automatic quantification with homemade Image J macros, allowing the study of more than 5,000 cells per condition in at least five independent donors, assessed these parameters.

### HIV-1 and cell-to-cell transfer

Freshly isolated CD14^+^ monocytes from HS transfected with siRNA against Siglec-1 or siRNA control were allowed to adhere in the absence of serum (2×10^6^ cells/6-well in 1.5 mL). After 4h of culture, RPMI-1640 supplemented with 20 ng/mL M-CSF and 20% FBS were added to the cells (vol/vol). After 24h, cells were conditioned with cmMTB media. At day 4, 120 ng p24 of a HIV-1 NLAD8 strain pseudotyped with a VSVG envelope was used to infect half of the cells, kept in culture for 2 more days. At day 6, before co-culture, uninfected cells were stained with CellTracker Green CMFDA Dye (Thermo Fisher Scientific). For mitochondria transfer, half of the macrophages were pre-stained with Green CellTracker, and the other half, uninfected, was stained with mitoTracker Deep-Red prior to co-culture. Briefly, cells were washed with PBS Mg^2+^/Ca^2+^ and stained for 30 min with 500 ng/mL CellTracker or mitoTracker, before washing with RPMI-1640 10% FBS. HIV-1^+^ or mitoTracker^+^ and CellTracker^+^ cells were then detached using accutase (Sigma) and co-cultured at a 1:1 ratio on glass coverslips in 24-well.

### Histological analyses

Paraffin embedded tissue samples were sectioned and stained with hematoxylin and eosin for histomorphological analysis. Different antigen unmasking methods where used on tissue slides for immunohistochemical staining, which was performed using anti-CD163 (Leica/Novocastra), anti-Siglec-1 (Novus Biologicals) and anti-pSTAT1 (Cell Signaling Technology). Sections were then incubated with biotin-conjugated polyclonal anti-mouse or anti-rabbit immunoglobulin antibodies followed by the streptavidin-biotin-peroxidase complex (ABC) method (Vector Laboratories). Finally, sections were counter-stained with hematoxylin. Slides were scanned with the Panoramic 250 Flash II (3DHISTECH). Virtual slides were automatically quantified for macrophage distribution as previously described (Souriant et al., 2019). Immunofluorescence staining of the sections was performed as described above and followed by anti-mouse IgG isotype specific or anti-rabbit IgG antibodies labelled with Alexa488 and Alexa555 (Molecular Probes). Samples were mounted with Prolong^®^ Antifade reagent (Molecular Probes) and examined using a 60×/1.40N.A. objective of an Olympus spinning disk microscope.

### Quantification and statistical analysis

Information on the statistical tests used and the exact values of n (donors) can be found in the Figure Legends. All statistical analyses were performed using GraphPad Prism 8.0.0 (GraphPad Software Inc.). Two-tailed paired or unpaired t-test was applied on data sets with a normal distribution (determined using Kolmogorov-Smirnov test), whereas two-tailed Mann-Whitney (unpaired test) or Wilcoxon matched-paired signed rank tests were used otherwise. p<0.05 was considered as the level of statistical significance (* p≤0.05; ** p≤0.005; *** p≤0.0005; **** p≤0.0001).

## ACKNOWLEDGMENTS

We greatly acknowledge F. Capilla and T. Al Saati, US006/CREFRE for histology analyses; P. Constant, F. Levillain, F. Moreau and C. Berrone, IPBS and Genotoul Anexplo-IPBS, for accessing the BSL3 facilities; E. Näser, E. Vega, A. Peixoto, S. Mazeres and the Genotoul TRI-IPBS facilities for imaging and flow cytometry. We thank F. Quiroga and C. del Carmen Melucci Ganzarain, Instituto de Investigaciones Biomédicas en Retrovirus y SIDA, INBIRS UBA - CONICET, Buenos Aires, Argentina, for the technical help and advice provided. We greatly thank Y-M. Boudehen for technical expertise provided in molecular biology, M. Dalod and B. Raynaud-Messina for fruitful discussions, and S. Benichou for providing HIV-1 strains. We are grateful to D. Hudrisier, C. Gutierrez, C. A. Spinner, L. Bernard, and B. Raymond for critical reading of the manuscript and helpful comments. This work was supported by the *Centre National de la Recherche Scientifique*, *Université Paul Sabatier*, the *Agence Nationale de la Recherche* (ANR14-CE11-0020-02, ANR16-CE13-0005-01, ANR-11-EQUIPEX-0003), the *Agence Nationale de Recherche sur le Sida et les hépatites virales* (ANRS2014-CI-2, ANRS2014-049, ANRS2018-01), the ECOS-Sud program (A14S01), the *Fondation pour la Recherche Médicale* (DEQ2016 0334894; DEQ2016 0334902), the *Fondation Bettencourt Schueller*, INSERM Plan Cancer, the Argentinean National Agency of Promotion of Science and Technology (PICT-2015-0055 and PICT-2017-1317). We also thank the AIDS Research and Reference Reagent Program, Division of AIDS, NIAID. The NHP study was supported by NIH award OD011104, AI058609, AI111943 and AI111914. The genetic analyses were realized within the framework of the Swiss HIV Cohort Study (SHCS Project number 717), which is supported by the Swiss National Science Foundation (Grant Number 148522) and by the SHCS research foundation. M.D. is supported by an ATP (*Axes Thématiques Prioritaires*) doctoral scholarship from *Université Paul Sabatier*, S.S. by a 4^th^-year doctoral scholarship from Sidaction, and S. R. by a scholarship from Toulouse University Hospital to perform a Master’s degree. J.M.-P and NI-U are supported by the Spanish Secretariat of State of Research, Development and Innovation through grant SAF2016-80033-R, J.M.-P. by the Spanish AIDS network *Red Temática Cooperativa de Investigación en SIDA*, and S.B. by the *Rio Hortega programme* funded by the Spanish Health Institute Carlos III (No. CM17/00242).

## MOVIE LEGENDS

**Movie 1** (Related to Figure 3A).

Z-stack of confocal microscopy images, showing Siglec-1 (red), microtubules (MT, green) and F-actin (grey) of day 6 HIV-1 infected macrophages, treated with cmMTB. Siglec-1 localizes on the thick MT+ F-actin+ TNT but not on thin MT-F-actin+ TNTs.

**Movie 2** (Related to Figure 3D).

3D reconstitution of confocal images, showing Siglec-1 (red), HIV-1 Gag (green) and WGA (grey) of day 6 HIV-1 infected macrophages, treated with cmMTB.

---

**Figure S1.**
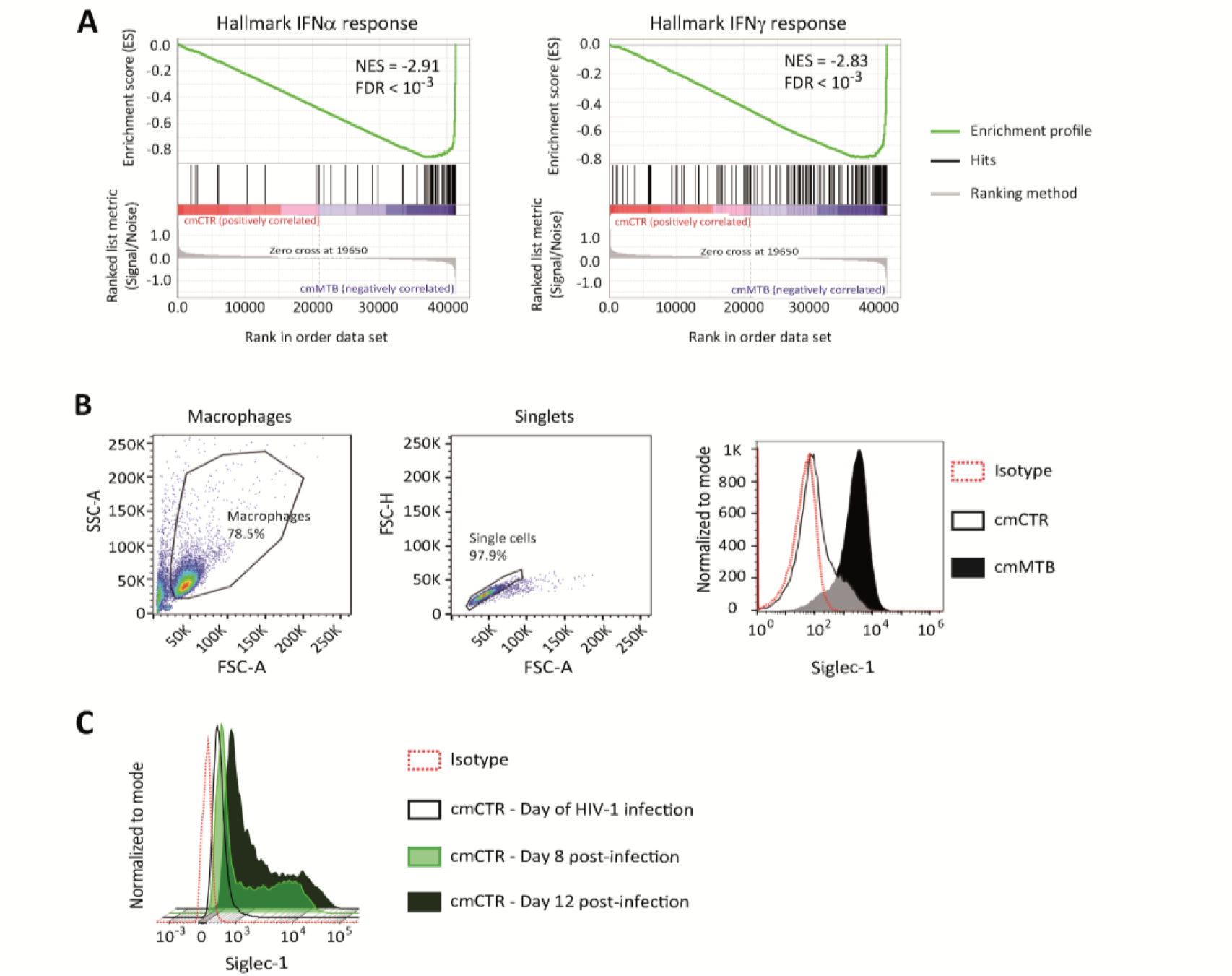
Tuberculosis-associated microenvironment increases Siglec-1 expression in human macrophages (see also Figure 1). (A-C) For 3 days, human monocytes were differentiated into macrophages with cmCTR (white) or cmMTB (black) supernatants. (A) (Left) Gene set enrichment plot of the interferon alpha (IFNα) response (hallmark collection of MSigDB). This plot shows the distribution of the barcode between macrophages exposed to cmCTR (red) *versus* cmMTB (blue) supernatants. Each bar of the barcode corresponds to a signature gene of the gene set. The skewing to the right indicates enrichment in macrophages exposed to cmMTB *versus* cmCTR supernatant of genes up-regulated in response to IFNα. (Right) Gene set enrichment plot of the IFNγ response (hallmark collection of MSigDB). (B-C) Flow cytometry gating strategy to assess Siglec-1 cell-surface expression in human macrophages exposed to cmCTR (white) and cmMTB (black) (B), or cmCTR-treated cells infected with HIV-1 (C). (B) Left: Based on size (FCS-A) and granularity (SSC-A), a gate was created to separate human macrophages from cell debris and dying cells. Macrophages were then subjected through a second gate based FSC Area Scaling (FCS-A and FCS-H) to separate singlets from doublets. Right: Based on the singlet gate, the histogram plot illustrates Siglec-1 expression that is higher in cmMTB- than in cmCTR-treated macrophages. (C) The histogram plot illustrates the time-course of Siglec-1 expression that is upregulated after HIV-1 infection in cmCTR-treated cells compared to uninfected ones.

**Figure S2.**
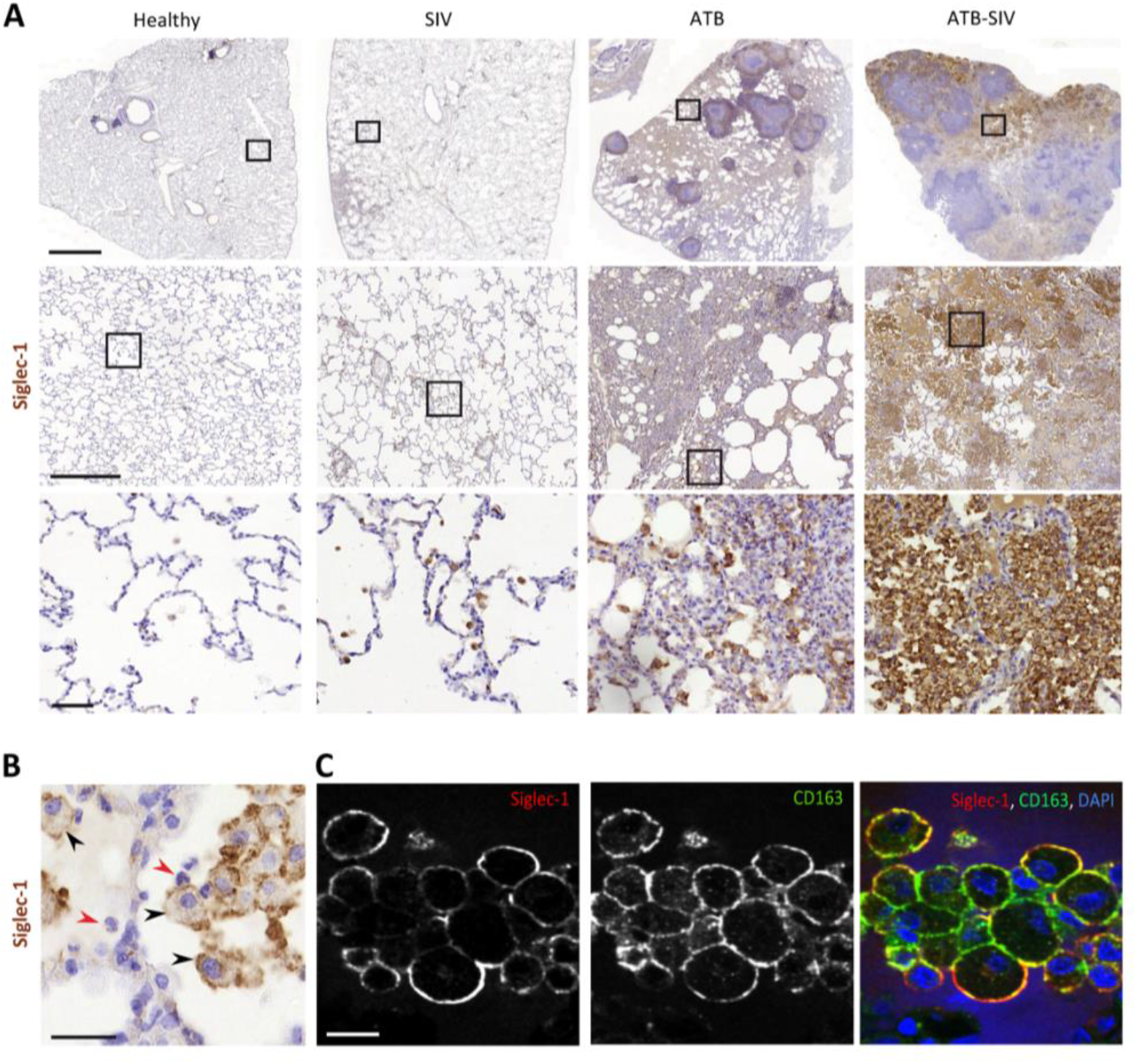
Tuberculosis-associated microenvironment increases Siglec-1 expression in non-human primate alveolar macrophages (see also Figure 1). (A) Accumulation of Siglec-1^+^ alveolar macrophages in the lung of co-infected non-human primates (NHP). Representative immunohistochemical images of Siglec-1 staining (brown) in lung biopsies of healthy, SIV-infected (SIV), active tuberculosis (ATB) and co-infected with SIV (ATB-SIV) NHP. Scale bars from top to bottom: 2 mm, 500 μm and 50 μm. (B-C) Siglec-1^+^ cells display the alveolar macrophage morphology. (B) Representative immunohistochemistry image from lung biopsy of an ATB-SIV NHP stained for Siglec-1 (brown). Siglec-1^+^ cells display a cell morphology with a single nucleus and large cytoplasm reminiscent of macrophage (black arrowhead); Siglec-1^−^ cells display a different nucleus morphology and small cytoplasm reminiscent of neutrophils (red arrowhead). Scale bar, 20 μm. (C) Representative immunofluorescence images of alveolar macrophages found in lung biopsy of a representative ATB-SIV NHP stained for Siglec-1 (red), CD163 (green) and DAPI (nuclei, blue). Scale bar, 20 μm.

**Figure S3.**
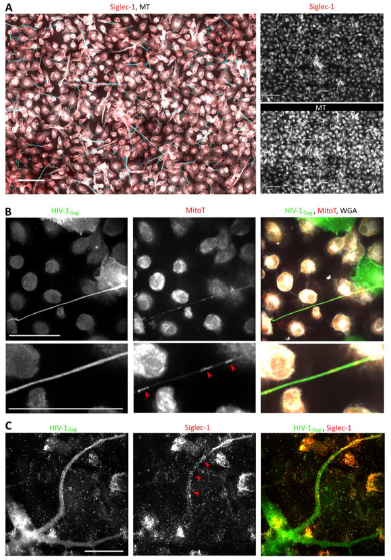
Siglec-1 localizes specifically on thick tunneling nanotubes that contain HIV-1 Gag and mitochondria (see also Figure 3). (A-C) Human monocytes were differentiated into macrophages with cmMTB for 3 days, infected with HIV-1-ADA strain (unless indicated otherwise) and then fixed at day 3 (A-B) or 10 (C) post-infection. (A) Representative immunofluorescence images used for semi-automatic quantification of TNT in cmMTB-treated macrophages infected with HIV-1. Cells were stained for extracellular Siglec-1 (red), intracellular tubulin (MT, grey) and Wheat Germ Agglutinin (WGA, not shown). Blue lines show all TNT considered. Thick (WGA^+^, MT^+^) and thin (WGA^+^, MT^−^) TNT were assessed for Siglec-1 positivity by applying a threshold and measured in length. Scale bar, 20 μm. (B) Representative immunofluorescence images of cmMTB-treated macrophages infected with HIV-1-NLAD8-VSVG, loaded with Mitotracker (MitoT, red) and stained for intracellular HIV-1 Gag (green) and WGA (grey). Red arrowheads show mitochondria inside HIV-1 Gag-containing TNT. Inserts are 2× zoom. Scale bar, 20 μm. (C) Representative immunofluorescence images of cmMTB-treated macrophages infected with HIV-1 and kept in culture until day 10 post-infection. Cells were fixed and stained for intracellular HIV-1 Gag (green), extracellular Siglec-1 (red) and WGA (grey). Red arrowheads show Siglec-1 on HIV-1 Gag-containing TNT emanating from an infected multinucleated giant cell (MGC). Scale bar, 20 μm.

**Figure S4.**
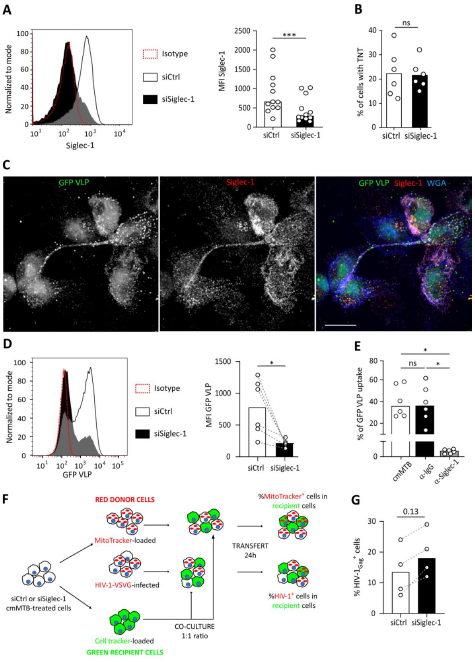
Siglec-1 is required for the capture and transfer of HIV-1 in cmMTB-treated macrophages (see also Figure 4). (A-D, F-G) Monocytes from healthy subjects were transfected with siRNA targeting of Siglec-1 (siSiglec-1) or not (siCtrl). A day after, monocytes were differentiated into macrophages with cmMTB for 3 days. (A) Representative histogram (left) and vertical scatter plot showing the median fluorescent intensity (MFI) (right) of Siglec-1 expression on the indicated cell populations. (B) Vertical scatter plot indicating the percentage of cells forming TNT in cells. (C-E) Inhibition of Siglec-1 reduces binding of HIV-1 Gag−eGFP VLP (GFP VLP). (C) Representative immunofluorescence (IF) images of cmMTB-treated cells incubated with GFP VLP (green) for 3.5h. Cells were fixed and stained for Siglec-1 (red) and Wheat Germ Agglutinin (WGA, blue). Scale bar, 500 μm. (D) Representative histogram (left) and vertical scatter plot showing the median fluorescent intensity (MFI) (right) displaying of GFP VLP binding in the indicated cell populations. (E) Vertical scatter plot showing the percentage of GFP VLP binding in cmMTB treated cells pre-incubated with specific anti-Siglec-1 (α-Siglec-1, grey), isotype control antibody (α-IgG, black) or mock (white). (E) Experimental model for the cell-to-cell transfer experiment. siRNA-transfected donor (red) cells were either labelled with MitoTracker (1h prior co-culture) or infected with HIV-1-NLAD8-VSVG (48h prior to co-culture), while autologous recipient cells (green) were stained with cell tracker (1h prior co-culture). Both donor and recipient cells were co-culture for 24h. Red fluorescence was then measured in recipient cells by IF. (G) Vertical scatter plot showing the percentage of HIV-1_Gag_^+^ cells at the time of co-culture experiment in the indicated cells. Statistical analyses: Two-tailed, Wilcoxon matched-pairs signed rank test (A, B, D-E, G). *P < 0.05, ***P < 0.001. ns: not significant.

## REFERENCES

Akiyama, H., Ramirez, N.G., Gudheti, M.V., and Gummuluru, S. (2015). CD169-mediated trafficking of HIV to plasma membrane invaginations in dendritic cells attenuates efficacy of anti-gp120 broadly neutralizing antibodies. PLoS pathogens 11, e1004751.

Akiyama, H., Ramirez, N.P., Gibson, G., Kline, C., Watkins, S., Ambrose, Z., and Gummuluru, S. (2017). Interferon-Inducible CD169/Siglec1 Attenuates Anti-HIV-1 Effects of Alpha Interferon. Journal of virology 91.

Bell, L.C.K., and Noursadeghi, M. (2018). Pathogenesis of HIV-1 and Mycobacterium tuberculosis co-infection. Nature reviews Microbiology 16, 80–90.

Berry, M.P., Graham, C.M., McNab, F.W., Xu, Z., Bloch, S.A., Oni, T., Wilkinson, K.A., Banchereau, R., Skinner, J., Wilkinson, R.J., et al. (2010). An interferon-inducible neutrophil-driven blood transcriptional signature in human tuberculosis. Nature 466, 973–977.

Bracq, L., Xie, M., Benichou, S., and Bouchet, J. (2018). Mechanisms for Cell-to-Cell Transmission of HIV-1. Frontiers in immunology 9, 260.

Cai, Y., Sugimoto, C., Liu, D.X., Midkiff, C.C., Alvarez, X., Lackner, A.A., Kim, W.K., Didier, E.S., and Kuroda, M.J. (2015). Increased monocyte turnover is associated with interstitial macrophage accumulation and pulmonary tissue damage in SIV-infected rhesus macaques. Journal of leukocyte biology 97, 1147–1153.

Cribbs, S.K., Lennox, J., Caliendo, A.M., Brown, L.A., and Guidot, D.M. (2015). Healthy HIV-1-infected individuals on highly active antiretroviral therapy harbor HIV-1 in their alveolar macrophages. AIDS research and human retroviruses 31, 64–70.

Crocker, P.R., Paulson, J.C., and Varki, A. (2007). Siglecs and their roles in the immune system. Nature reviews Immunology 7, 255–266.

Deffur, A., Mulder, N.J., and Wilkinson, R.J. (2013). Co-infection with Mycobacterium tuberculosis and human immunodeficiency virus: an overview and motivation for systems approaches. Pathogens and disease 69, 101–113.

Diedrich, C.R., and Flynn, J.L. (2011). HIV-1/mycobacterium tuberculosis coinfection immunology: how does HIV-1 exacerbate tuberculosis? Infection and immunity 79, 1407–1417.

Dupont, M., Souriant, S., Lugo-Villarino, G., Maridonneau-Parini, I., and Verollet, C. (2018). Tunneling Nanotubes: Intimate Communication between Myeloid Cells. Frontiers in immunology 9, 43.

Esmail, H., Riou, C., Bruyn, E.D., Lai, R.P., Harley, Y.X.R., Meintjes, G., Wilkinson, K.A., and Wilkinson, R.J. (2018). The Immune Response to Mycobacterium tuberculosis in HIV-1-Coinfected Persons. Annual review of immunology 36, 603–638.

Eugenin, E.A., Gaskill, P.J., and Berman, J.W. (2009). Tunneling nanotubes (TNT) are induced by HIV-infection of macrophages: a potential mechanism for intercellular HIV trafficking. Cellular immunology 254, 142–148.

Foreman, T.W., Mehra, S., LoBato, D.N., Malek, A., Alvarez, X., Golden, N.A., Bucsan, A.N., Didier, P.J., Doyle-Meyers, L.A., Russell-Lodrigue, K.E., et al. (2016). CD4+ T-cell-independent mechanisms suppress reactivation of latent tuberculosis in a macaque model of HIV coinfection. Proceedings of the National Academy of Sciences of the United States of America 113, E5636–5644.

Ganor, Y., Real, F., Sennepin, A., Dutertre, C.A., Prevedel, L., Xu, L., Tudor, D., Charmeteau, B., Couedel-Courteille, A., Marion, S., et al. (2019). HIV-1 reservoirs in urethral macrophages of patients under suppressive antiretroviral therapy. Nature microbiology 4, 633–644.

Gummuluru, S., Pina Ramirez, N.G., and Akiyama, H. (2014). CD169-dependent cell-associated HIV-1 transmission: a driver of virus dissemination. The Journal of infectious diseases 210 Suppl 3, S641–647.

Halasz, H., Ghadaksaz, A.R., Madarasz, T., Huber, K., Harami, G., Toth, E.A., Osteikoetxea-Molnar, A., Kovacs, M., Balogi, Z., Nyitrai, M., et al. (2018). Live cell superresolution-structured illumination microscopy imaging analysis of the intercellular transport of microvesicles and costimulatory proteins via nanotubes between immune cells. Methods and applications in fluorescence 6, 045005.

Hartnell, A., Steel, J., Turley, H., Jones, M., Jackson, D.G., and Crocker, P.R. (2001). Characterization of human sialoadhesin, a sialic acid binding receptor expressed by resident and inflammatory macrophage populations. Blood 97, 288–296.

Hashimoto, M., Bhuyan, F., Hiyoshi, M., Noyori, O., Nasser, H., Miyazaki, M., Saito, T., Kondoh, Y., Osada, H., Kimura, S., et al. (2016). Potential Role of the Formation of Tunneling Nanotubes in HIV-1 Spread in Macrophages. Journal of immunology 196, 1832–1841.

Honeycutt, J.B., Thayer, W.O., Baker, C.E., Ribeiro, R.M., Lada, S.M., Cao, Y., Cleary, R.A., Hudgens, M.G., Richman, D.D., and Garcia, J.V. (2017). HIV persistence in tissue macrophages of humanized myeloid-only mice during antiretroviral therapy. Nature medicine 23, 638–643.

Honeycutt, J.B., Wahl, A., Baker, C., Spagnuolo, R.A., Foster, J., Zakharova, O., Wietgrefe, S., Caro-Vegas, C., Madden, V., Sharpe, G., et al. (2016). Macrophages sustain HIV replication in vivo independently of T cells. The Journal of clinical investigation 126, 1353–1366.

Ivashkiv, L.B., and Donlin, L.T. (2014). Regulation of type I interferon responses. Nature reviews Immunology 14, 36–49.

Izquierdo-Useros, N., Lorizate, M., Contreras, F.X., Rodriguez-Plata, M.T., Glass, B., Erkizia, I., Prado, J.G., Casas, J., Fabrias, G., Krausslich, H.G., et al. (2012a). Sialyllactose in viral membrane gangliosides is a novel molecular recognition pattern for mature dendritic cell capture of HIV-1. PLoS biology 10, e1001315.

Izquierdo-Useros, N., Lorizate, M., McLaren, P.J., Telenti, A., Krausslich, H.G., and Martinez-Picado, J. (2014). HIV-1 capture and transmission by dendritic cells: the role of viral glycolipids and the cellular receptor Siglec-1. PLoS pathogens 10, e1004146.

Izquierdo-Useros, N., Lorizate, M., Puertas, M.C., Rodriguez-Plata, M.T., Zangger, N., Erikson, E., Pino, M., Erkizia, I., Glass, B., Clotet, B., et al. (2012b). Siglec-1 is a novel dendritic cell receptor that mediates HIV-1 trans-infection through recognition of viral membrane gangliosides. PLoS biology 10, e1001448.

Jambo, K.C., Banda, D.H., Kankwatira, A.M., Sukumar, N., Allain, T.J., Heyderman, R.S., Russell, D.G., and Mwandumba, H.C. (2014). Small alveolar macrophages are infected preferentially by HIV and exhibit impaired phagocytic function. Mucosal immunology 7, 1116–1126.

Jaroenpool, J., Rogers, K.A., Pattanapanyasat, K., Villinger, F., Onlamoon, N., Crocker, P.R., and Ansari, A.A. (2007). Differences in the constitutive and SIV infection induced expression of Siglecs by hematopoietic cells from non-human primates. Cellular immunology 250, 91–104.

Kabaso, D., Lokar, M., Kralj-Iglic, V., Veranic, P., and Iglic, A. (2011). Temperature and cholera toxin B are factors that influence formation of membrane nanotubes in RT4 and T24 urothelial cancer cell lines. International journal of nanomedicine 6, 495–509.

Kaushal, D., Mehra, S., Didier, P.J., and Lackner, A.A. (2012). The non-human primate model of tuberculosis. Journal of medical primatology 41, 191–201.

Kuroda, M.J., Sugimoto, C., Cai, Y., Merino, K.M., Mehra, S., Arainga, M., Roy, C.J., Midkiff, C.C., Alvarez, X., Didier, E.S., et al. (2018). High Turnover of Tissue Macrophages Contributes to Tuberculosis Reactivation in Simian Immunodeficiency Virus-Infected Rhesus Macaques. The Journal of infectious diseases 217, 1865–1874.

Lastrucci, C., Benard, A., Balboa, L., Pingris, K., Souriant, S., Poincloux, R., Al Saati, T., Rasolofo, V., Gonzalez-Montaner, P., Inwentarz, S., et al. (2015). Tuberculosis is associated with expansion of a motile, permissive and immunomodulatory CD16(+) monocyte population via the IL-10/STAT3 axis. Cell research 25, 1333–1351.

Lokar, M., Kabaso, D., Resnik, N., Sepcic, K., Kralj-Iglic, V., Veranic, P., Zorec, R., and Iglic, A. (2012). The role of cholesterol-sphingomyelin membrane nanodomains in the stability of intercellular membrane nanotubes. International journal of nanomedicine 7, 1891–1902.

Martinez-Picado, J., McLaren, P.J., Telenti, A., and Izquierdo-Useros, N. (2017). Retroviruses As Myeloid Cell Riders: What Natural Human Siglec-1 “Knockouts” Tell Us About Pathogenesis. Frontiers in immunology 8, 1593.

Mathews, S., Branch Woods, A., Katano, I., Makarov, E., Thomas, M.B., Gendelman, H.E., Poluektova, L.Y., Ito, M., and Gorantla, S. (2019). Human Interleukin-34 facilitates microglia-like cell differentiation and persistent HIV-1 infection in humanized mice. Molecular neurodegeneration 14, 12.

Mattila, J.T. (2019). Type 1 interferon expression and signaling occur in spatially-distinct regions in granulomas from Mycobacterium tuberculosis-infected cynomolgus macaques. The Journal of Immunology 202, 62.19–62.19.

McNab, F., Mayer-Barber, K., Sher, A., Wack, A., and O’Garra, A. (2015). Type I interferons in infectious disease. Nature reviews Immunology 15, 87–103.

Mehra, S., Golden, N.A., Dutta, N.K., Midkiff, C.C., Alvarez, X., Doyle, L.A., Asher, M., Russell-Lodrigue, K., Monjure, C., Roy, C.J., et al. (2011). Reactivation of latent tuberculosis in rhesus macaques by coinfection with simian immunodeficiency virus. Journal of medical primatology 40, 233–243.

Moreira-Teixeira, L., Mayer-Barber, K., Sher, A., and O’Garra, A. (2018). Type I interferons in tuberculosis: Foe and occasionally friend. The Journal of experimental medicine 215, 1273–1285.

O’Garra, A., Redford, P.S., McNab, F.W., Bloom, C.I., Wilkinson, R.J., and Berry, M.P. (2013). The immune response in tuberculosis. Annual review of immunology 31, 475–527.

O’Neill, A.S., van den Berg, T.K., and Mullen, G.E. (2013). Sialoadhesin - a macrophage-restricted marker of immunoregulation and inflammation. Immunology 138, 198–207.

Onfelt, B., Nedvetzki, S., Benninger, R.K., Purbhoo, M.A., Sowinski, S., Hume, A.N., Seabra, M.C., Neil, M.A., French, P.M., and Davis, D.M. (2006). Structurally distinct membrane nanotubes between human macrophages support long-distance vesicular traffic or surfing of bacteria. Journal of immunology 177, 8476–8483.

Osteikoetxea-Molnar, A., Szabo-Meleg, E., Toth, E.A., Oszvald, A., Izsepi, E., Kremlitzka, M., Biri, B., Nyitray, L., Bozo, T., Nemeth, P., et al. (2016). The growth determinants and transport properties of tunneling nanotube networks between B lymphocytes. Cellular and molecular life sciences: CMLS 73, 4531–4545.

Pino, M., Erkizia, I., Benet, S., Erikson, E., Fernandez-Figueras, M.T., Guerrero, D., Dalmau, J., Ouchi, D., Rausell, A., Ciuffi, A., et al. (2015). HIV-1 immune activation induces Siglec-1 expression and enhances viral trans-infection in blood and tissue myeloid cells. Retrovirology 12, 37.

Puryear, W.B., Akiyama, H., Geer, S.D., Ramirez, N.P., Yu, X., Reinhard, B.M., and Gummuluru, S. (2013). Interferon-inducible mechanism of dendritic cell-mediated HIV-1 dissemination is dependent on Siglec-1/CD169. PLoS pathogens 9, e1003291.

Puryear, W.B., Yu, X., Ramirez, N.P., Reinhard, B.M., and Gummuluru, S. (2012). HIV-1 incorporation of host-cell-derived glycosphingolipid GM3 allows for capture by mature dendritic cells. Proceedings of the National Academy of Sciences of the United States of America 109, 7475–7480.

Redondo-Morata, L., Giannotti, M.I., and Sanz, F. (2012). Influence of cholesterol on the phase transition of lipid bilayers: a temperature-controlled force spectroscopy study. Langmuir: the ACS journal of surfaces and colloids 28, 12851–12860.

Rempel, H., Calosing, C., Sun, B., and Pulliam, L. (2008). Sialoadhesin expressed on IFN-induced monocytes binds HIV-1 and enhances infectivity. PLoS One 3, e1967.

Ritchie, M.E., Phipson, B., Wu, D., Hu, Y., Law, C.W., Shi, W., and Smyth, G.K. (2015). limma powers differential expression analyses for RNA-sequencing and microarray studies. Nucleic acids research 43, e47.

Rodrigues, V., Ruffin, N., San-Roman, M., and Benaroch, P. (2017). Myeloid Cell Interaction with HIV: A Complex Relationship. Frontiers in immunology 8, 1698.

Rustom, A., Saffrich, R., Markovic, I., Walther, P., and Gerdes, H.H. (2004). Nanotubular highways for intercellular organelle transport. Science 303, 1007–1010.

Sattentau, Q.J., and Stevenson, M. (2016). Macrophages and HIV-1: An Unhealthy Constellation. Cell host & microbe 19, 304–310.

Schneider, W.M., Chevillotte, M.D., and Rice, C.M. (2014). Interferon-stimulated genes: a complex web of host defenses. Annual review of immunology 32, 513–545.

Sewald, X., Ladinsky, M.S., Uchil, P.D., Beloor, J., Pi, R., Herrmann, C., Motamedi, N., Murooka, T.T., Brehm, M.A., Greiner, D.L., et al. (2015). Retroviruses use CD169-mediated trans-infection of permissive lymphocytes to establish infection. Science 350, 563–567.

Souriant, S., Balboa, L., Dupont, M., Pingris, K., Kviatcovsky, D., Cougoule, C., Lastrucci, C., Bah, A., Gasser, R., Poincloux, R., et al. (2019). Tuberculosis Exacerbates HIV-1 Infection through IL-10/STAT3-Dependent Tunneling Nanotube Formation in Macrophages. Cell reports 26, 3586–3599 e3587.

Subramanian, A., Tamayo, P., Mootha, V.K., Mukherjee, S., Ebert, B.L., Gillette, M.A., Paulovich, A., Pomeroy, S.L., Golub, T.R., Lander, E.S., et al. (2005). Gene set enrichment analysis: a knowledge-based approach for interpreting genome-wide expression profiles. Proceedings of the National Academy of Sciences of the United States of America 102, 15545–15550.

Thayanithy, V., Babatunde, V., Dickson, E.L., Wong, P., Oh, S., Ke, X., Barlas, A., Fujisawa, S., Romin, Y., Moreira, A.L., et al. (2014). Tumor exosomes induce tunneling nanotubes in lipid raft-enriched regions of human mesothelioma cells. Experimental cell research 323, 178–188.

Torralba, D., Baixauli, F., and Sanchez-Madrid, F. (2016). Mitochondria Know No Boundaries: Mechanisms and Functions of Intercellular Mitochondrial Transfer. Frontiers in cell and developmental biology 4, 107.

Toth, E.A., Oszvald, A., Peter, M., Balogh, G., Osteikoetxea-Molnar, A., Bozo, T., Szabo-Meleg, E., Nyitrai, M., Derenyi, I., Kellermayer, M., et al. (2017). Nanotubes connecting B lymphocytes: High impact of differentiation-dependent lipid composition on their growth and mechanics. Biochimica et biophysica acta Molecular and cell biology of lipids 1862, 991–1000.

Troegeler, A., Lastrucci, C., Duval, C., Tanne, A., Cougoule, C., Maridonneau-Parini, I., Neyrolles, O., and Lugo-Villarino, G. (2014). An efficient siRNA-mediated gene silencing in primary human monocytes, dendritic cells and macrophages. Immunology and cell biology 92, 699–708.

VanderVen, B.C., Huang, L., Rohde, K.H., and Russell, D.G. (2016). The Minimal Unit of Infection: Mycobacterium tuberculosis in the Macrophage. Microbiology spectrum 4.

Verollet, C., Souriant, S., Bonnaud, E., Jolicoeur, P., Raynaud-Messina, B., Kinnaer, C., Fourquaux, I., Imle, A., Benichou, S., Fackler, O.T., et al. (2015). HIV-1 reprograms the migration of macrophages. Blood 125, 1611–1622.

Verollet, C., Zhang, Y.M., Le Cabec, V., Mazzolini, J., Charriere, G., Labrousse, A., Bouchet, J., Medina, I., Biessen, E., Niedergang, F., et al. (2010). HIV-1 Nef triggers macrophage fusion in a p61Hck- and protease-dependent manner. Journal of immunology 184, 7030–7039.

Yu, Y.R., Hotten, D.F., Malakhau, Y., Volker, E., Ghio, A.J., Noble, P.W., Kraft, M., Hollingsworth, J.W., Gunn, M.D., and Tighe, R.M. (2016). Flow Cytometric Analysis of Myeloid Cells in Human Blood, Bronchoalveolar Lavage, and Lung Tissues. American journal of respiratory cell and molecular biology 54, 13–24.

Ziegler-Heitbrock, L., Lotzerich, M., Schaefer, A., Werner, T., Frankenberger, M., and Benkhart, E. (2003). IFN-alpha induces the human IL-10 gene by recruiting both IFN regulatory factor 1 and Stat3. Journal of immunology 171, 285–290.

Zou, Z., Chastain, A., Moir, S., Ford, J., Trandem, K., Martinelli, E., Cicala, C., Crocker, P., Arthos, J., and Sun, P.D. (2011). Siglecs facilitate HIV-1 infection of macrophages through adhesion with viral sialic acids. PLoS One 6, e24559.

